# Coordinated multiple cellular processes in tongue development

**DOI:** 10.1101/2023.01.16.524234

**Authors:** Maiko Kawasaki, Katsushige Kawasaki, Finsa Tisna Sari, Takehisa Kudo, Jun Nihara, Madoka Kitamura, Takahiro Nagai, Vanessa Utama, Yoko Ishida, Fumiya Meguro, Takayuki Nishimura, Yuan Kogure, Satoshi Maruyama, Jun-ichi Tanuma, Yoshito Kakihara, Takeyasu Maeda, Sarah Ghafoor, Roman H. Khonsari, Pierre Corre, Paul T. Sharpe, Martyn T. Cobourne, Brunella Franco, Atsushi Ohazama

## Abstract

Dysfunction of primary cilia leads to genetic disorder, ciliopathies, which shows various malformations in many vital organs such as brain. Multiple tongue deformities including cleft, hamartoma and ankyloglossia are also seen in ciliopathies, which yield difficulties in fundamental functions such as mastication and vocalization. Here, we found these tongue anomalies in mice with mutation of ciliary protein. Abnormal cranial neural crest-derived cells (CNCC) failed to evoke Hh signal for differentiation of mesoderm-derived cells into myoblasts, which resulted in abnormal differentiation of mesoderm-derived cells into adipocytes. The ectopic adipose subsequently arrested migration of other mesoderm-derived cells and CNCC. Some aberrant CNCC abnormally differentiated into osteoblasts due to the lack of Hh signal, which migrated into tongue to form ectopic bone. Ankyloglossia was caused by aberrant cell migration due to lack of non-canonical Wnt signaling. In addition to ciliopathies, these tongue anomalies are often observed as non-familial condition in human. We found that these tongue deformities could be reproduced in wild-type mice by simple mechanical manipulations in CNCC to disturb cellular processes which were disrupted in mutant mice. Thus, tongue development requires coordinated multiple cellular processes (cell-cell contact, migration and differentiation). Our results provide hints for possible future treatment in ciliopathies.

## Introduction

The primary cilium, a nonmotile organelle existing on almost all somatic cell surfaces in vertebrates, has been shown to play critical roles in many biological processes, including regulating signaling pathways (Evans et al., 2006, Bisgrove and Yost 2006). The primary cilium consists of a membrane-bound cylinder surrounding nine doublet microtubules, which extends from a centriole termed the basal body. Cilia are assembled and maintained by intraflagellar transport (IFT), in which multiple protein complexes are moved bidirectionally along the axoneme. Perturbations in the function of primary cilia result in a wide spectrum of human diseases: the ciliopathies (Rosenbaum and Witman, 2002; Pan et al., 2005; Bisgrove and Yost. 2006; Evans et al., 2006; Scholey and Anderson, 2006). Ciliopathy patients exhibit prominent mixed symptoms in several vital organs including the brain, lung, kidney and liver, and others, such as the eye and digit. Multiple congenital tongue anomalies including aglossia, cleft, hamartoma and ankyloglossia are also observed in ciliopathy patients (Mostafa et al., 2005, Auber et al., 2007, Zaghloul & Brugmann 2011). These tongue anomalies significantly impair quality of life, since the tongue plays a critical role in multiple fundamental functions including mastication, deglutition, general sensation and taste, oral cleansing and vocalization. In addition to ciliopathies, these tongue anomalies are often observed as non-familial condition (Cobourne et al., 2019, Hill et al., 2020, Li et al., 2020, Yin et al., 2020). However, the cause of these tongue anomalies remains unclear.

The tongue is a muscular organ. During tongue development, tongue myoblasts originate from the anterior-most somites, which are derived from the mesoderm (Parada et al., 2012, Ziermann et al., 2018, Sambasivan et al., 2011). Cells from these somites migrate to the craniofacial region via bilateral pathways, known collectively as the hypoglossal cord (Sambasivan et al., 2011). Cranial neural crest-derived cells (CNCC) also migrate into the tongue primordia, and contribute to form the lamina propria, tendon and interstitial connective tissue to compartmentalize tongue muscles and serve as their attachments. A lingual swelling begin to appear in the mandibular processes at embryonic day (E) 11.5 in mice, which undergoes rapid enlargement, with a rudimentary tongue-like structure observed by E12.5.

OFDI syndrome is an X-linked dominant ciliopathy that affects females and causes prenatal lethality in males (Macca & Franco 2009). OFD1 syndrome is characterized by malformations of the oroface and digits (Macca and Franco 2009). *OFD1* has been identified as the gene mutated in OFD1 syndrome patients. The *OFD1* gene is located on the X-chromosome and the encoded protein is localized to basal bodies of primary cilia. Congenital tongue anomalies are also observed in OFD1 syndrome patients (Mostafa et al., 2005, Auber et al., 2007, Zaghloul & Brugmann 2011).

Here, we describe that tongue development require proper multiple cellular processes, and genetic or mechanical disruption of multiple cellular processes in CNCC attribute to congenital tongue malformations as familial and non-familial conditions, respectively. Our findings contribute to understand coordinated cellular processes during organogenesis.

## Results

### Tongue phenotypes in Ofd1 mutant

We firstly examined mice with conditional deletion of *Ofd1* in CNCC using *Wnt1Cre*. Both hemizygous [*Ofd1^fl^;Wnt1Cre(HM)*] and heterozygous *Ofd1* mutant mice [*Ofd1^fl/WT^;Wnt1Cre(HET)*] die at birth. *Ofd1^fl^;Wnt1Cre(HM)* mice have aglossia (Fig. 1B; Okuhara et al., 2019), suggesting that complete deletion of *Ofd1* in CNCC leads to a failure of tongue formation. In contrast, *Ofd1^fl/WT^;Wnt1Cre(HET)* mice have an obvious abnormal-shaped tongue, characterized by the presence of clefts and multiple protrusions (Fig. 1D, S1; n=58/58). These morphological anomalies varied widely in *Ofd1^fl/WT^;Wnt1Cre(HET)* mice.

**Figure 1.**
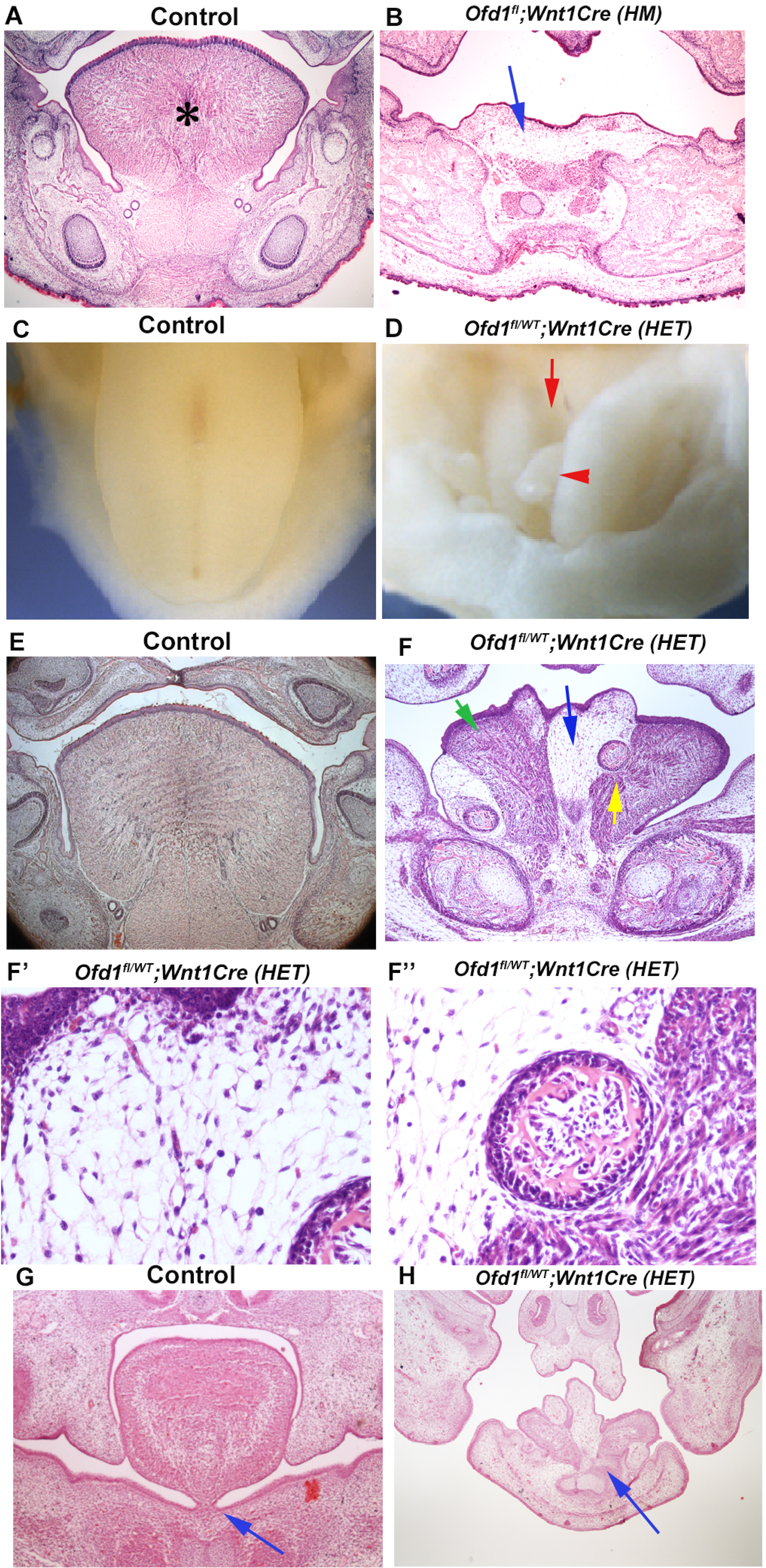
Tongue phenotypes in *Ofd1* mutant mice. (A, B, E-H) Frontal sections showing histological images in wild-type (A, E, G), *Ofd1^fl^;Wnt1Cre(HM)* (B) and *Ofd1^fl/WT^;Wnt1Cre(HET)* (F-F’’, H) at E18.5. Arrow indicating sparse tissue (B). *: tongue (A). Green, blue and yellow arrows indicating normal muscle, ectopic sparse tissue and ectopic bone, respectively (F). F’ and F’’ are high magnification of F indicated by blue and yellow arrow, respectively. Arrows indicating tongue frenum region (G, H). (C, D) Image showing oral view of tongue in wild-type (C) and *Ofd1^fl/WT^;Wnt1Cre(HET)* (D). Arrowhead and arrow indicating ectopic protrusion and cleft, respectively (D).

Ectopic bone and sparse tissue were observed in the tongue of *Ofd1^fl/WT^;Wnt1Cre(HET)* mice, although normal muscle was also present (Fig. 1F-1F’’). The size and location of ectopic bone and sparse tissue varied widely in these mice (Fig. S2). Protrusions consistted of sparse tissue. A lack of tongue frenum was often observed in *Ofd1^fl/WT^;Wnt1Cre(HET)* mice (n=30/58; Fig. 1H).

To identify the type of sparse tissue found in *Ofd1^fl/WT^;Wnt1Cre(HET)* mice, we analyzed gene expression following laser microdissection. Results of PCR and immunohistochemistry analysis suggested that this tissue was brown adipose tissue, but not muscle or white adipose tissue (Fig. 2A-2C, S3). Protrusion was thus hamartoma of adipose tissue. Although mucous salivary glands associated-adipose and intermuscular adipose are known to be present in wild-type tongue, we found that the brown adipose tissue found in the *Ofd1* mutant tongue was not related to these adipose (see Fig. S4).

**Figure 2.**
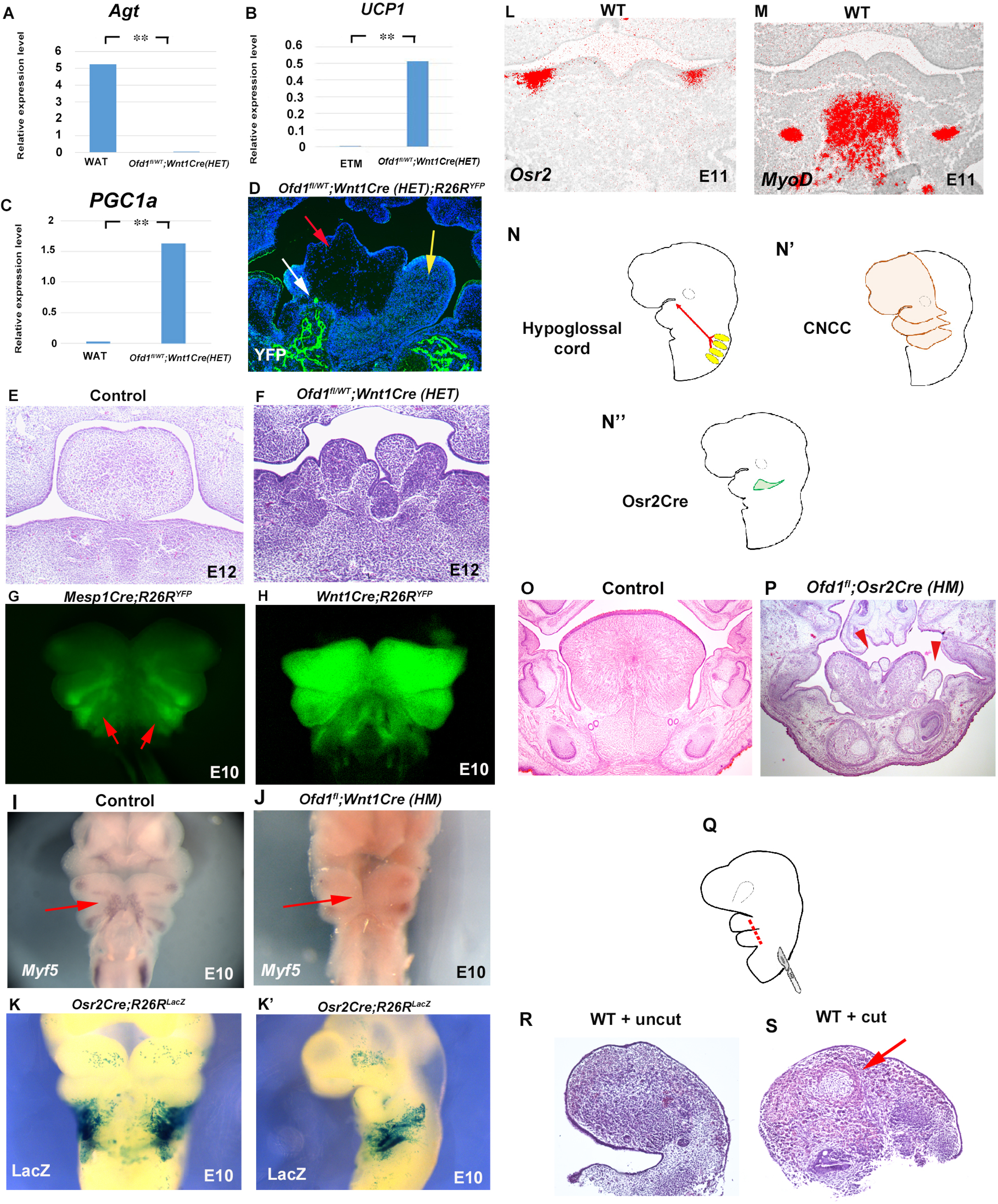
Sparse tissue in *Ofd1* mutant tongue. (A-C) q-PCR on mRNA isolated from ectopic sparse tissue, white adipose tissue (WAT) and embryonic tongue muscle (ETM). ** P<0.01. Only low-level expression of myogenic markers in the sparse tissue compared with the embryonic tongue muscle (A). Markers for brown adipocytes were expressed at high-level in the sparse tissue, in comparison with those in the embryonic tongue muscle (B) or adult white adipose tissue (C). (D-F) Frontal sections showing immunohistochemistry of YFP (D) and histological images (E, F) in *Ofd1^fl/WT^;Wnt1Cre(HET);R26R^YFP^* (D), wild-type (E) and *Ofd1^fl/WT^;Wnt1Cre(HET)* (F) at E12.5 (E, F) and E16.5 (D). Red, yellow and white arrow indicating that sparse, muscle and bone tissue, respectively (D). (G, H, I, J) Frontal images of *YFP* expression (G, H) and *Myf5* expression of whole mount *in situ* hybridization (I, J) in *Mesp1Cre;R26R^YFP^* (G), *Wnt1Cre;R26R^YFP^* (H), wild-type (I) and *Ofd1^fl^;Wnt1Cre(HM)* (J). (K, K’) Frontal (K) and sagittal (K’) view of LacZ stained *Osr2Cre;R26R^LacZ^* mice. (L, M) Frontal sections showing *in situ* hybridization of *Osr2* (L) and *MyoD* (M) in wild-type. (N-N’’) Schematic diagram showing hypoglossal cord (arrow in N), occipital somites (yellow in N), CNCC (orange in N’) and *Osr2Cre* expression domain (green in N’’). (O, P) Frontal sections showing histological images in wild-type (O) and *Ofd1^fl^;Osr2Cre(HM)* (P) at E18.5. Arrowheads indicating sparse tissue. (Q) Scheme diagram showing lateral view of craniofacial region with incision (red line). (R, S) Histological images of cultured wild-type tongue without incision (R) and with incision (S). Arrowheads indicating ectopic sparse tissue (P).

These anomalies could not be detected in mice with epithelial conditional deletion of *Ofd1* [*Ofd1^fl/WT^;K14Cre(HET)* or *Ofd1^fl^;K14Cre(HM)*]

### Cell differentiation by cell-cell interaction

Brown adipose tissue is normally derived from the mesoderm (Kajimura et al., 2010), whereas *Ofd1* is deleted in CNCC in *Ofd1^fl/WT^;Wnt1Cre(HET)* mice. In order to understand whether ectopic brown adipose tissue in the *Ofd1* mutant tongue was derived from CNCC, we examined YFP expression in *Ofd1^fl/WT^;Wnt1Cre(HET);R26R^YFP^* mice. No YFP expression could be detected in the adipose tissue, indicating that ectopic brown adipose was derived from the mesoderm (Fig. 2D). Tongue muscles are also derived from the mesoderm in wild-type mice, and tongue muscle in *Ofd1^fl/WT^;Wnt1Cre(HET);R26R^YFP^* mice exhibited no YFP expression, suggesting that muscular tissue in mutant tongue was derived from the mesoderm under normal conditions (Fig. 2D). In contrast, bone in the tongue of *Ofd1^fl/WT^;Wnt1Cre(HET)* mice was found to be derived from CNCC, since it showed YFP expression (Fig. 2D). Thus, in the mutant tongue, the brown adipose and muscle tissue were derived from the mesoderm, whilst ectopic bone was from CNCC. Ectopic brown adipose tissue was formed from mesoderm-derived cells, when *Ofd1* was deleted from CNCC. It is possible that the interaction between CNCC and mesoderm-derived cells is involved in differentiation of mesoderm-derived cells. To investigate this hypothesis, we firstly examined the early stages of tongue development to identify the timing and location of possible interactions between CNCC and mesoderm-derived cells. Morphological anomalies were observed in the tongue of *Ofd1^fl/WT^;Wnt1Cre(HET)* mice at E12.5, when the early tongue swelling has become apparent (Fig. 2F), indicating that mesoderm-derived cells were already demonstrating an abnormal status when they migrated to form the tongue swelling. Before mesoderm-derived cells enter the mandibular processes, they migrate as a single stream along the hypoglossal cord around E9-E10. We confirmed the presence of mesoderm-derived cells and CNCC in the wild-type hypoglossal cord at E10 using *Mesp1Cre;R26R^YFP^* and *Wnt1Cre;R26R^YFP^* mice, respectively (Fig. 2G, 2H). In addition, *Myf5* expression was observed in the hypoglossal cord region of wild-type mice at the time, which could not be detected when *Ofd1* was deleted from all CNCC [*Ofd1^fl^;Wnt1Cre(HM)*; Fig. 2I, 2J]. Thus, mesoderm-derived cells failed to differentiate into myoblast before they entered to mandibular processes in *Ofd1* mutant mice. In other word, in wild-type mice, mesoderm-derived cells start to differentiate into myoblasts within the hypoglossal cord. In wild-type mice, *Osr2* expression partially overlapped with the hypoglossal cord and CNCC at E10 (Fig. 2K, 2K’). At E11, both *Osr2*-expressing cells and mesoderm-derived cells were present in the mandibular process; however, *Osr2*-expressing cells did not interact with mesoderm-derived cells at the stage, due to the significant distance between *Osr2*-expressing cells and mesoderm-derived cells (confirmed by *MyoD* or *Myf5*) (Fig. 2L, 2M, S5). *Osr2*-expressing cells likely interact with mesoderm-derived cells only within the hypoglossal cord at E10 (Fig. 2N-2N”). Therefore, we generated *Ofd1* mutant mice using *Osr2Cre* to identify whether mesoderm-derived cells is related to CNCC within the hypoglossal cord for their proper differentiation. Indeed, ectopic sparse tissue was observed in *Ofd1^fl^;Osr2Cre(HM)* mice (Fig. 2P, Supple info. 1). To further confirm whether differentiation of mesoderm-derived cells is relied on interaction between mesoderm-derived cells and CNCC within the hypoglossal cord, we made several incisions into the hypoglossal cord in wild-type mice to reduce the interaction, and then cultured the embryos. To minimize disturbance of their migration, punctiform incisions were made under the visualization of mesoderm-derived cells using *Mesp1Cre;R26R^YFP^* mice as wild-type mice (Fig. 2G, 2Q). Adipose-like tissue was observed in the tongue of wild-type with incisions (n=4/8, Fig. 2R), which was never seen in wild-type without incisions (n=0/8, Fig. 2S). These results indicated that in normal tongue development, CNCC interact with mesoderm-derived cells within the hypoglossal cord, which is essential for normal differentiation of mesoderm-derived cells into myoblasts. Aberrant interaction results in abnormal differentiation of mesoderm-derived cells into brown adipocytes. Thus, ectopic adipose formation due to aberrant differentiation of mesoderm-derived cells were caused by a lack of proper cell-cell interaction between CNCC and mesoderm-derived cells.

In addition to ectopic bone and adipose tissues, normal muscle tissue was also present in *Ofd1^fl/WT^;Wnt1Cre(HET)* tongue (see Fig. 1F), indicating that some CNCC successfully interact and induced normal differentiation of mesoderm-derived cells into muscle progenitors, whilst other CNCC populations failed to elicit this. Thus, two types of CNCC (normal and abnormal CNCC) would appear to be present in *Ofd1^fl/WT^;Wnt1Cre(HET)* mice. This is most likely caused by X-inactivation, since *Ofd1* is located on the X-chromosome. To confirm this, we used *Hprt^GFP^* mice that have a GFP reporter expressed by Cre ribonuclease on the X-chromosome. Normal CNCC (CNCC with *Ofd1*) could be seen as GFP*-*expressing cells in *Ofd1^fl/WT^;Wnt1Cre(HET);Hprt^GFP^* mice, when these mice were generated through crossing female *Ofd1^fl/fl^* mice with male *Wnt1Cre;Hprt^YFP^* mice. YFP*-*expressing cells were observed in some region of the mandibular processes in *Ofd1^fl/WT^;Wnt1Cre(HET);Hprt^GFP^* mice, indicating that normal CNCC were present in *Ofd1^fl/WT^;Wnt1Cre(HET)* mice (Fig. 3A). In control mice, Myo-D was observed as a single domain (Fig. 3B); however, *Ofd1^fl/WT^;Wnt1Cre(HET)* mice showed a mosaic pattern of MyoD-positive domains (Fig. 3C) as YFP domain in *Ofd1^fl/WT^;Wnt1Cre(HET);Hprt^GFP^* mice showed (Fig. 3A). Furthermore, MyoD-negative domains were found around GFP*-*negative domains in hypoglossary cord of *Ofd1^fl/WT^;Wnt1Cre(HET);Hprt^GFP^* mice (Fig. 3D, 3E; Fig. 3D and 3E were obtained from same mutant mouse). These findings confirmed that, in *Ofd1^fl/WT^;Wnt1Cre(HET)* mice, CNCC displaying inactivation of the X-chromosome with *Ofd1* mutation induced normal differentiation of surrounding mesoderm-derived cells, while CNCC with inactivation of the normal X-chromosome failed to elicit this to surrounding mesoderm-derived cells. It is likely that CNCC in the hypoglossal cord are randomly divided into two types of cell clusters (cells with and cells without *Ofd1*) due to the X-inactivation, since X-inactivation is based on random choice between the two X-chromosomes. This might lead to the phenotypic tongue variation seen in *Ofd1^fl/WT^;Wnt1Cre(HET)* mice. Thus, the interaction between CNCC and mesoderm-derived cells is under control of X-inactivation.

**Figure 3.**
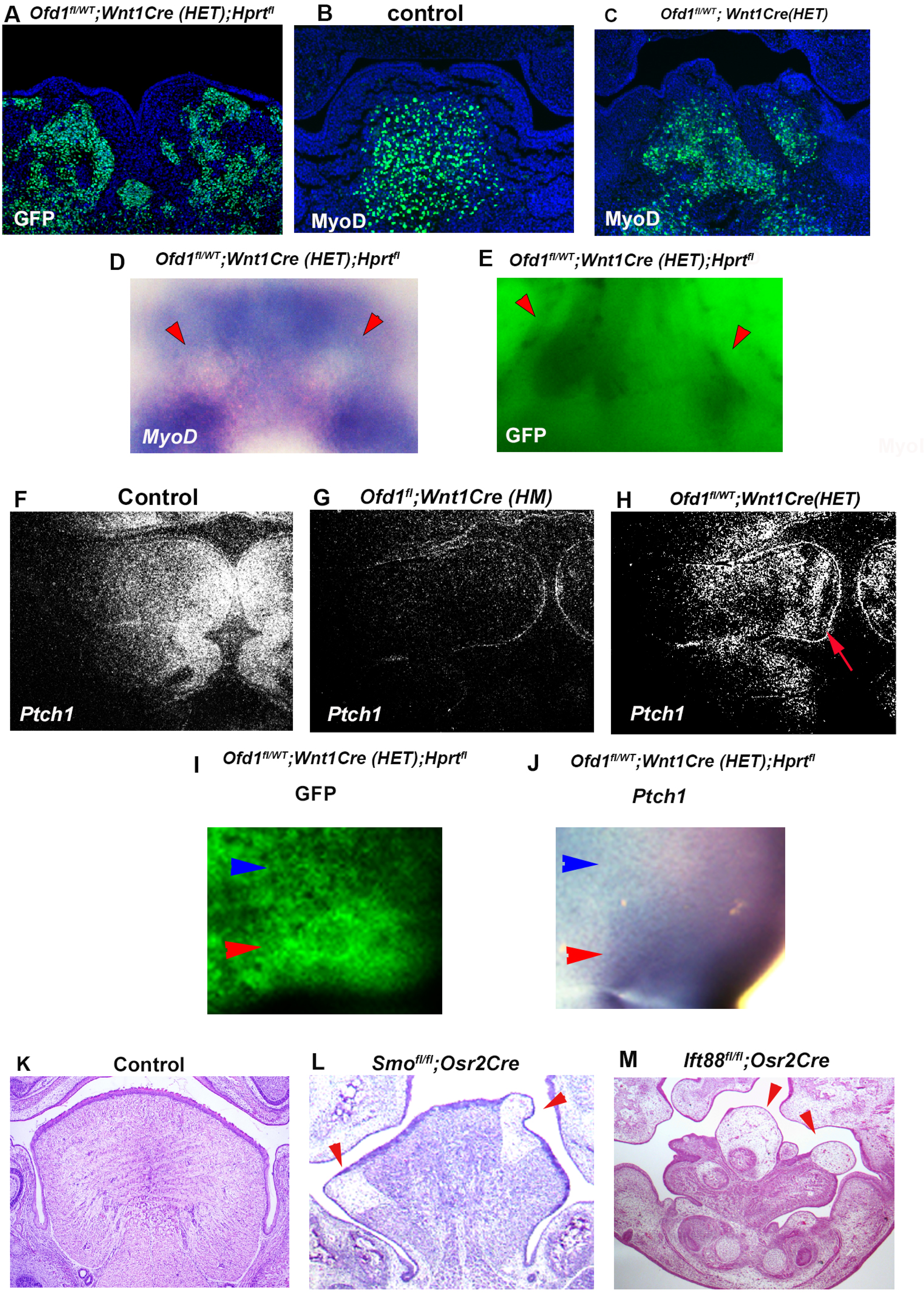
Hh signal in *Ofd*1 mutant tongue. (A-C, F-H) Frontal sections showing immunohistochemistry of GFP (A), MyoD (B, C) and *in situ* hybridization of *Ptch1* (F-H) in *Ofd1^fl/WT^;Wnt1Cre(HET);Hprt^fl^* (A), wild-type (B, F), *Ofd1^fl^;Wnt1Cre(HM)* (G) and *Ofd1^fl/WT^;Wnt1Cre(HET)* (C, H) mice. Arrow indicating mosaic *Ptch1* expression (H). (D, E) Frontal view showing *Gfp* expression (E) before fixation (E) and whole mount *in situ* hybridization of *MyoD* (D) after fixation in same *Ofd1^fl/WT^;Wnt1Cre(HET);Hprt^fl^* mouse. Arrowheads indicating MyoD-(D) and Gfp- (E) negative domains. (I, J) Frontal view showing *Gfp* expression before fixation (I) and whole mount *in situ* hybridization of *Ptch1* after fixation (J) in same *Ofd1^fl/WT^;Wnt1Cre(HET);Hprt^fl^* mouse. Blue arrows indicating Ptch1- (J) and Gfp- (I) negative domains. Red arrows indicating Ptch1- (J) and Gfp- (I) positive domains. (K-M) Frontal sections showing histology of wild-type (K), *Smo^fl/fl^;Osr2Cre* (L) and *Ift88^fl/fl^;Osr2Cre* (N) at E18.5. Arrowheads indicating ectopic sparse tissue.

*Ofd1* mutation in CNCC disrupts proper interactions between CNCC and mesoderm-derived cells, and subsequent differentiation of mesoderm-derived cells, suggesting that signalling from CNCC is likely to be crucial for these interaction and subsequent differentiation. *Ptch1* (major mediators of Hh signaling) was expressed in the wild-type hypoglossal cord, but not in the *Ofd1^fl^;Wnt1Cre(HM)* cord, suggesting that *Ofd1* mutation from all CNCC led to a complete absence of Hh signal activation in the hypoglossal cord (Fig. 3F, 3G). On the other hand, in the hypoglossal cord of *Ofd1^fl/WT^;Wnt1Cre(HET)* mice, a mosaic pattern of *Ptch1*-negative and -positive domains were observed, which varied between *Ofd1^fl/WT^;Wnt1Cre(HET)* mice (Fig. 3H). These are also probably caused by X-inactivation. In fact, *Ptch1*-negative and -positive domains were overlapped with GFP*-*negative and -positive domains in *Ofd1^fl/WT^;Wnt1Cre(HET);Hprt^GFP^* mice, respectively (Fig. 3I, 3J). To understand whether ectopic brown adipose tissue formation was due to downregulation of the Hh signaling pathway in CNCC within the hypoglossal cord, we generated mice with conditional deletion of *Smo* (an essential mediator of Hh signaling activity) using *Osr2Cre* (*Smo^fl/fl^;Osr2Cre*) mice. Adipose-like tissue was observed in the tongue of *Smo^fl/fl^;Osr2Cre* mice (Fig. 3L). Thus, the interaction between CNCC and mesoderm-derived cells are reliant on Hh signalling in CNCC, which is essential for proper differentiation of mesoderm-derived cells. Hh signaling is activated in primary cilia, and tongue anomalies are observed in ciliopathy patients. We also found alteration of primary cilia in CNCC of *Ofd1^fl^;Wnt1Cre(HM);R26R^YFP^* mice (Fig. S6). To confirm whether the disruption of primary cilia function induce tongue anomalies, we generated and examined mice with conditional deletion of *Ift88* (another primary cilia protein) using *Osr2Cre* (*Ift88^fl/fl^;Osr2Cre*). *Ift88^fl/fl^;Osr2Cre* mice also exhibited same abnormal tongue as those in *Ofd1^fl/WT^;Wnt1Cre(HET)* mice (Fig. 2M).

### Cell migration for tongue swelling

*Ofd1^fl^;Wnt1Cre(HM)* mice exhibit aglossia, indicating that an early tongue swelling does not occur when *Ofd1* is deleted from all CNCC (Fig. 4B, Okuhara et al., 2019). Before the emergence of a tongue swelling in wild-type mice (at E11.5), both CNCC and mesoderm-derived cells are present in the mandibular processes where tongue swelling emerge (Fig. 4C, 4D). However, *Ofd1^fl^;Wnt1Cre(HM);R26R^Yfp^* mice show no CNCC in the region (Fig. 4E), suggesting that CNCC with *Ofd1* mutation fail to migrate into the region. Moreover, no myogenic progenitors were present in this region (Fig. 4F), confirming that differentiation of mesoderm-derived cells into myoprogenitors does not occur in *Ofd1^fl^;Wnt1Cre(HM)* mice due to a lack of *Ofd1* in all CNCC (see Fig. 2J). On the other hand, we did find sparse-like tissue in this region of *Ofd1^fl^;Wnt1Cre(HM)* mice at this time (Fig. 4H). At later stages, the sparse tissue became apparent in the region where the tongue usually emerged (Fig. 4J and see Fig. 4B). PCR analysis showed that the sparse tissue found in *Ofd1^fl^;Wnt1Cre(HM)* mice was also brown adipose (Fig. 4K, 4L, S7A-S7C). We already concluded that ectopic brown adipose formation was caused by the lack of Hh signaling in CNCC. Indeed, the ectopic brown adipose was also observed in same region of mice with *Smo* deletion from all CNCC (*Smo^fl/fl^;Wnt1Cre* mice; Fig. S7E, S7G-S7P). These results confirmed that abnormal brown adipocytes derived from the mesoderm succeeded in migrating into the mandibular processes in *Ofd1^fl^;Wnt1Cre(HM)* mice, although CNCC with *Ofd1* mutation failed to migrate into this region. Thus, the interaction between CNCC and mesoderm-derived cells in the hypoglossal cord affect subsequent cell migration.

**Figure 4.**
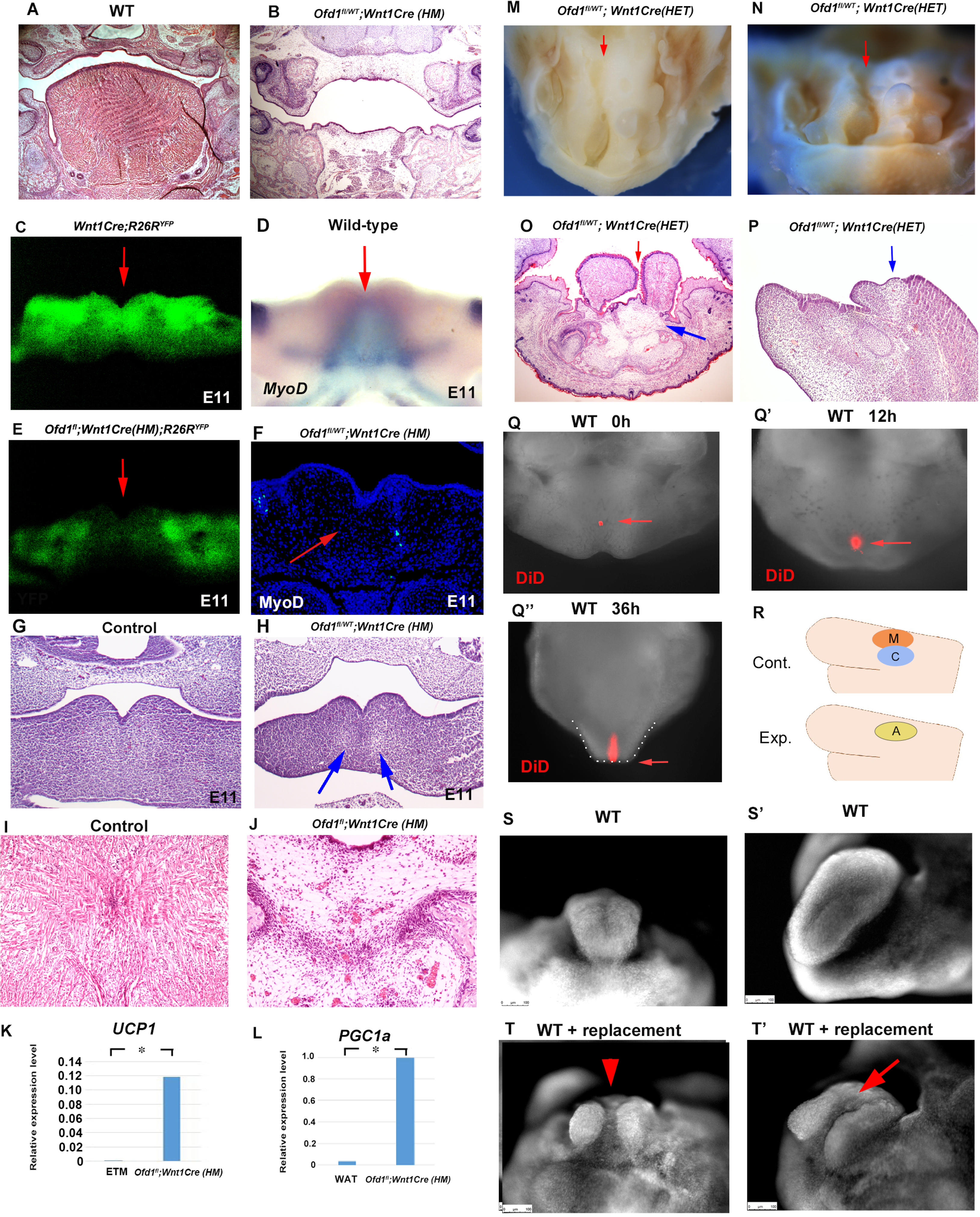
Migration of adipocytes in *Ofd1* mutants. (A, B, G-J, O, P) Frontal (A, B, G-J, O) and sagittal (P) sections showing histological images. Blue and red arrows indicating sparse tissue and cleft, respectively (H, O, P). (C, D, E) Frontal images of YFP expression (C, E) and whole mount *in situ* hybridization of *MyoD* (D) in *Wnt1Cre;R26R^YFP^* (C), wild-type (D) and *Ofd1^fl^;Wnt1Cre(HM);R26R^YFP^* (E). Arrows indicating region tongue swelling emerge. (F) Frontal section showing immunohistochemistry of MyoD in *Ofd1^fl^;Wnt1Cre(HM)*. Arrow indicating region tongue swelling emerge. (K, L) q-PCR on mRNA isolated from ectopic sparse tissue from *Ofd1^fl^;Wnt1Cre(HM)* mice, embryonic tongue muscle (ETM) and white adipose tissue (WAT). ** P<0.01. (M, N) Oral (M) and frontal (N) view of *Ofd1^fl/WT^;Wnt1Cre(HET)* mice. Arrows indicating cleft. (Q-Q’’) Nuclear fluorescent image showing oral view of wild-type mandible with DiD at E11.5 before culture (Q), 12h (Q’) and 36h (Q’’) after culture. Arrow indicating DiD labeled cells. Tongue like structure is outlined by white dots (Q’’). (R) Scheme diagram showing sagittal view of mandible exhibiting replacement of CNCC (C; blue circle) and myoblasts (M; red circle) into adipose tissue (A; yellow circle). (S-T’) Nuclear fluorescent image showing frontal (S, T) and oral (S’, T’) view of wild-type mandible without (S, S’) and with (T, T’) replacement.

The absence of a tongue swelling in *Ofd1^fl^;Wnt1Cre(HM)* mice suggested that ectopic brown adipocytes could not develop a tongue swelling, although they could migrate into mandibular processes. Unlike *Ofd1^fl^;Wnt1Cre(HM)* mice, tongue was present in *Ofd1^fl/WT^;Wnt1Cre(HET)* mice, and abnormal deep clefts originating from the tongue root were often observed, while adipose tissue was seen at the bottom of these clefts (Fig. 4M-4O, S1). Sagittal sections of the cleft part of the tongue also showed that the adipose tissue was present at the most posterior part of the cleft (Fig. 4P). Fate mapping analysis using DiD labelling (injection where the early tongue emerges=TE region; Fig. 4Q) confirmed that the tongue develops along the antero-positerior axis (Fig. 4Q-4Q’’). This suggested the possibility that the presence of adipose disturb the formation of the early tongue swelling, which lead to the cleft formation in *Ofd1^fl/WT^;Wnt1Cre(HET)* mice, since ectopic adipose could not form tongue swelling in *Ofd1^fl^;Wnt1Cre(HM)* mice. To confirm this hypothesis, myoblasts and CNCC were replaced with adipose tissue in wild-type mandible at E11.0 where the tongue emerges, and cultured them (Fig. 4R). The replacement was performed only restricted area within the region tongue swelling usually emerges. The cleft-like structures were observed in the cultured tongue, which was never seen in cultured tongue without replacement (Fig. 4S-4T’). Thus, cleft is occurred when adipose tissue was present which is surrounded by CNCC/myoblasts. These also suggested that tongue morphology was determined by which tissue first reaches the region where the tongue swelling emerges. Tongue swelling occur when CNCC/myoblasts reach first, while it is inhibited when adipocytes reach first.

We already concluded that ectopic adipose tissue could not form a tongue swelling; however, adipose tissue was often seen within the tongue swelling of *Ofd1^fl/WT^;Wnt1Cre(HET)* mice (Fig. 5A, 5B). Our findings indicated that early formation of the tongue swelling requires the presence of CNCC and mesoderm-derived cells. We often found that adipose tissue was present in the dorsum region of the emerging tongue swelling at E12.5, when the tongue swelling emerged in *Ofd1^fl/WT^;Wnt1Cre(HET)* mice (Fig. 5C, 5C’). It is possible that myogenic progenitors and CNCC push adipocytes up during early formation of the tongue swelling. To confirm this, we examined the formation of tongue papillae. In wild-type mice, the tongue papillae initiate through epithelial-mesenchymal interactions during emergence of the early tongue swelling. It is unlikely that adipocytes induce tongue papillae epithelium through epithelial-mesenchymal interaction. Therefore, the presence or absence of tongue papillae would be an indicator of whether adipocytes are located underneath the tongue epithelium during emergence of the tongue swelling. The epithelium and lamina propria overlying the tongue muscle exhibited tongue papillae-like structures in *Ofd1^fl/WT^;Wnt1Cre(HET)* mice (Fig. 5E, 5E’). However, only a thin epithelium was observed in the region overlying the ectopic adipose tissue (Fig. 5E, 5E’’). The tongue papillae marker *Krt1-5* (Jonker et al., 2004), was expressed in the epithelium overlying muscle tissue with lamina propria only, and not in the overlying ectopic adipose tissue (Fig. 5G). Thus, the presence of ectopic adipose tissue underneath the tongue epithelium led to a lack of tongue papillae. To further confirm this finding, we implanted adipose tissue on CNCC and myoblasts in wild-type mandibular explants at E11.0, and cultured them (Fig. 5H). The adipose tissue was found on the dorsum region after tongue swelling (Fig. 5I, 5J). Thus, adipocytes could enter the tongue swelling when adipocytes are located above CNCC and/or myoblasts. These also confirmed that tongue morphology was determined by which tissue first reaches the region where the tongue swelling emerges. Adipose could enter to tongue primordia when adipocytes reach after CNCC/myoblasts. Adipose tissue often goes down to the floor of the mandible from the dorsum of the tongue during emergence of tongue swelling (Fig. 5L; see 5A and 5B).

**Figure 5.**
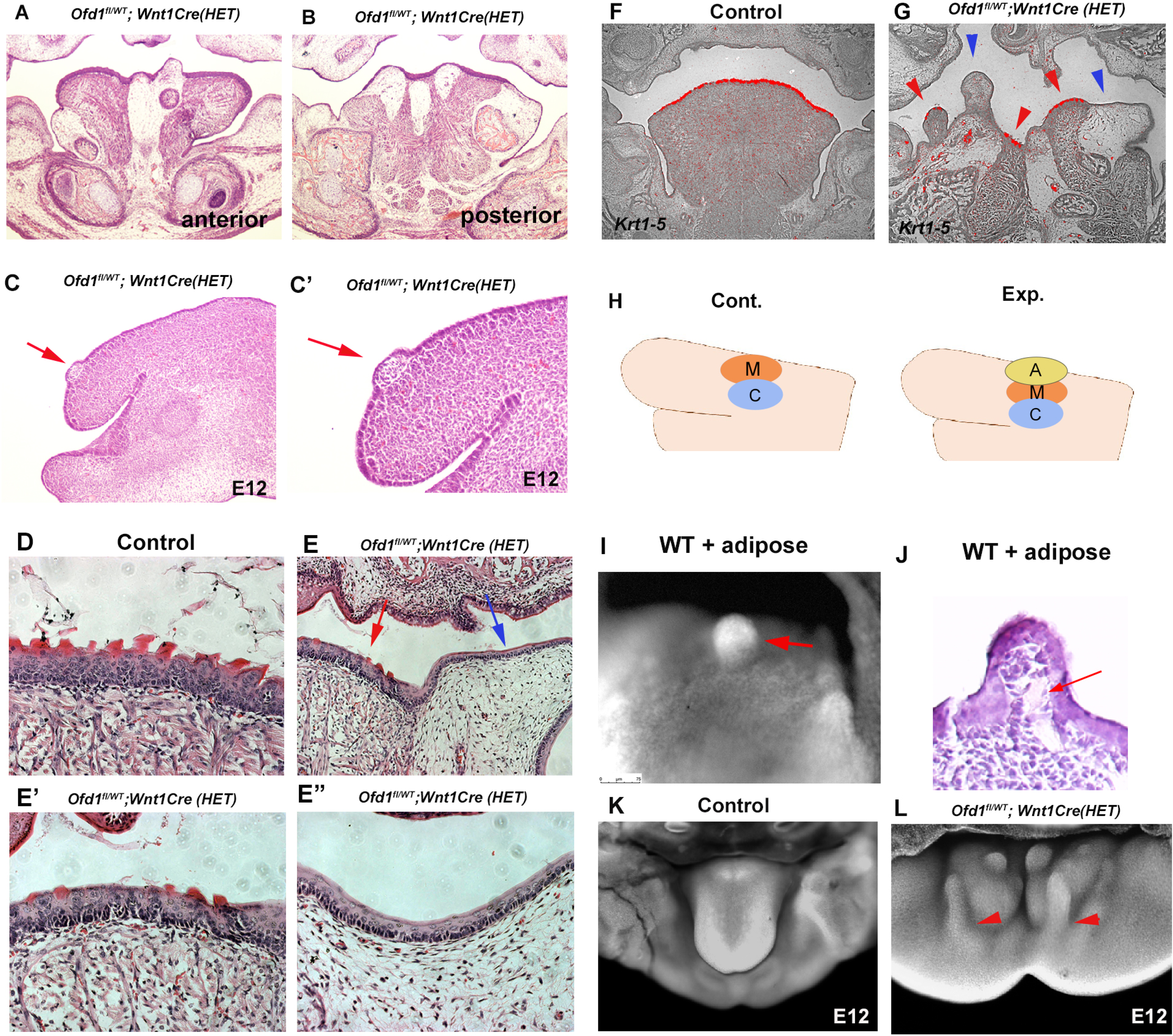
Adipose tissue in *Ofd1* mutant tongue. (A, B, C-E’’) Frontal (A, B, D-E’’) and sagittal (C, C’) sections showing histological images in wild-type (D), and *Ofd1^fl/WT^;Wnt1Cre(HET)* (A-C’, E-E’’). E’ and E’’ are high magnification of E. (F, G) Frontal sections showing *in situ* hybridization of *Krt1-5* in wild-type (F) and *Ofd1^fl/WT^;Wnt1Cre(HET)* (G). (H) Scheme diagram showing sagittal view of mandible exhibiting graft of adipose tissue (A; yellow circle), CNCC (C; blue circle) and myoblasts (M; red circle). (I) Nuclear fluorescent image showing dorsum of tongue in cultured mandible with grafting. (J) Histological images of dorsum of tongue in cultured mandible with grafting. (K, L) Nuclear fluorescent image showing oral view of developing mandibles in wild-type (K) and *Ofd1^fl/WT^;Wnt1Cre(HET)* (L). Arrowheads indicating protrusion into the floor of mandible from the dorsum surface of tongue swelling (L).

### Ectopic adipose in OFD1 patient

We next examined the tongue of an individual with *OFD1* G138S missense mutation. Several protrusions were observed in the tongue, which displayed an abnormally smooth surface in some regions, whilst others exhibited a normal dorsum surface (Fig. 6A, 6B). Similar to *Ofd1^fl/WT^;Wnt1Cre(HET)* mice, sparse tissue was observed in the tongue (Fig. 6E, 6F). PCR analysis (comparing cultured human skeletal muscle myoblasts or white adipocytes) indicated that the sparse tissue observed in the tongue of the OFD1 subject was brown adipose (Fig. 6G, 6H, S7Q-S7T). Furthermore, no obvious tongue papillae were detected in the protruded tissue (Fig. 6J), and expression of *KRT1-5* could not be detected in epithelium of the protruded tissue (Fig. 6L). Thus, our findings in mice likely occur in patient tongue formation.

**Figure 6.**
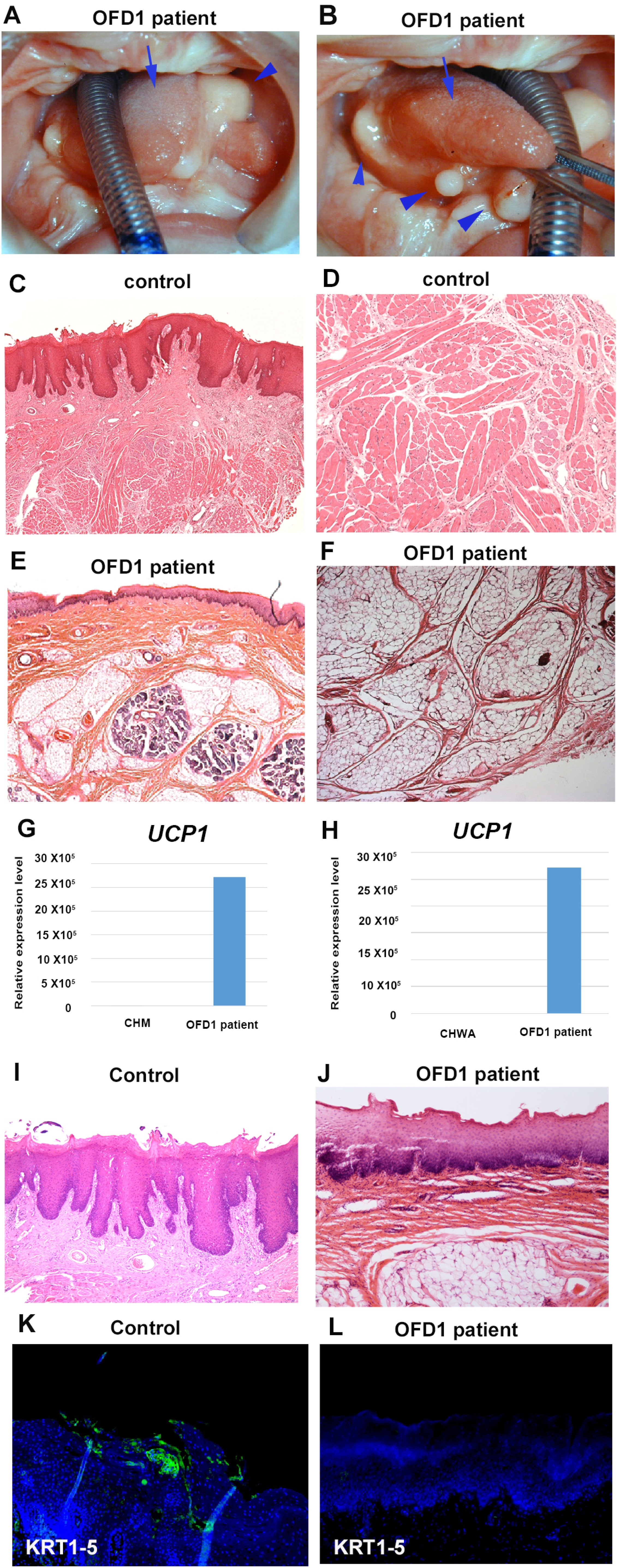
Tongue in OFD1 patient. (A, B) Tongue in OFD1 patient (G138S missense mutation). Arrowheads and arrows indicating protruded tissue and normal tongue, respectively. (C-F, I-L) Sections showing histological images (C-F, I, J) and immunohistochemistry of KRT1-5 (K, L) on control human tongue (C, D, I, K) and OFD1 patient tongue (E, F, J, L). (G, H) q-PCR on mRNA isolated from ectopic sparse tissue from OFD1 patient tongue, cultured human skeletal muscle myoblasts (CHM) and cultured human white adipocyte (CHWA).

### Cell differentiation and migration from mandibular bone region into developing tongue

Ectopic bone was observed in the tongue of *Ofd1^fl/WT^;Wnt1Cre(HET)* mice (see Fig. 1F). Ectopic bone formation in the tongue has been reported in human (Gorini et al., 2014). We already concluded that the ectopic bone seen in *Ofd1* mutant tongue is formed by CNCC (see Fig. 2D). In mandibular processes of *Ofd1^fl^;Wnt1Cre(HM)* mice, unlike region for emergence of early tongue swelling, CNCC with *Ofd1* mutation could migrate into the mandibular bone region, although all CNCC have no *Ofd1* (Fig. 7B). Thus, some CNCC retained the capability of migration in *Ofd1^fl^;Wnt1Cre(HM)* mice. We found the extra mandibular bone and Meckel’s cartilage formation lingual to the endogenous mandibular bone in *Ofd1^fl^;Wnt1Cre(HM)* mice, while these mice have no tongue (Fig. 7D, 7F). Hh signaling was downregulated in the lingual region (Fig. 7G, 7H). It has been shown that a lack of Hh signaling induce the duplication of the mandible at lingual region (Xu et al., 2019, Kitamura et al., 2020). These indicated that duplication of mandible was also occurred in *Ofd1^fl^;Wnt1Cre(HM)* mice due to the lack of Hh signaling. Unlike *Ofd1^fl^;Wnt1Cre(HM)* mice, tongue was present in *Ofd1^fl/WT^;Wnt1Cre(HET)* mice. Three-dimensional reconstruction in *Ofd1^fl/WT^;Wnt1Cre(HET)* mice showed that ectopic bone in the tongue was connected to the mandibular bone region (Fig. 7J). Ectopic osteoblasts was observed in early tongue swelling of *Ofd1^fl/WT^;Wnt1Cre(HET)* mice when tongue swelling emerged (Fig. 7L). The duplicated mandible remained along endogenous alveolar bone, when the tongue swelling was severely impaired in *Ofd1^fl/WT^;Wnt1Cre(HET)* mice (Fig. 7M). These suggested that ectopic tongue bone is likely to be consequence of migration of duplicated mandible into the tongue during emergence of tongue swelling. Ectopic bone was also observed in the tongue of *Ofd1* mutant mice using *Osr2Cre* [*Ofd1^fl^;Osr2Cre(HM)*], and it was found to be formed by *Osr2Cre*-expressing cells (Fig. 7N, 7O). In wild-type mice before emergence of tongue swelling (E11.0), *Osr2Cre* activation was observed in the lingual region where mandible duplicated, but not where endogenous mandibular bone formed (Fig. 7P, Lan et al., 2007). *Osr2Cre*-expressing cells are CNCC, since mesoderm-derived cells were not present in the lingual region of wild-type mice at the time (see Fig. 2L, 2M). After emergence of tongue swelling (E11.5), *Osr2* expression was also found within the tongue swelling in wild-type mice (Fig. 7Q, Lan et al., 2007). *Osr2*-expressing cells are likely to migrate into the tongue to form tongue connective tissue such as the lamina propria, tendon and interstitial connective tissue, since these tissues are formed by CNCC. Thus, cells in the lingual region usually differentiate into fibroblasts and migrate into tongue primordia to form tongue connective tissue. However, in *Ofd1^fl/WT^;Wnt1Cre(HET)* mice, these cells abnormally differentiate into osteoblasts due to the lack of Hh signaling, which migrate into the tongue when tongue formed. The abnormal differentiation occurs only when normal X-chromosome is inactivated in the lingual region (Fig. S8).

**Figure 7.**
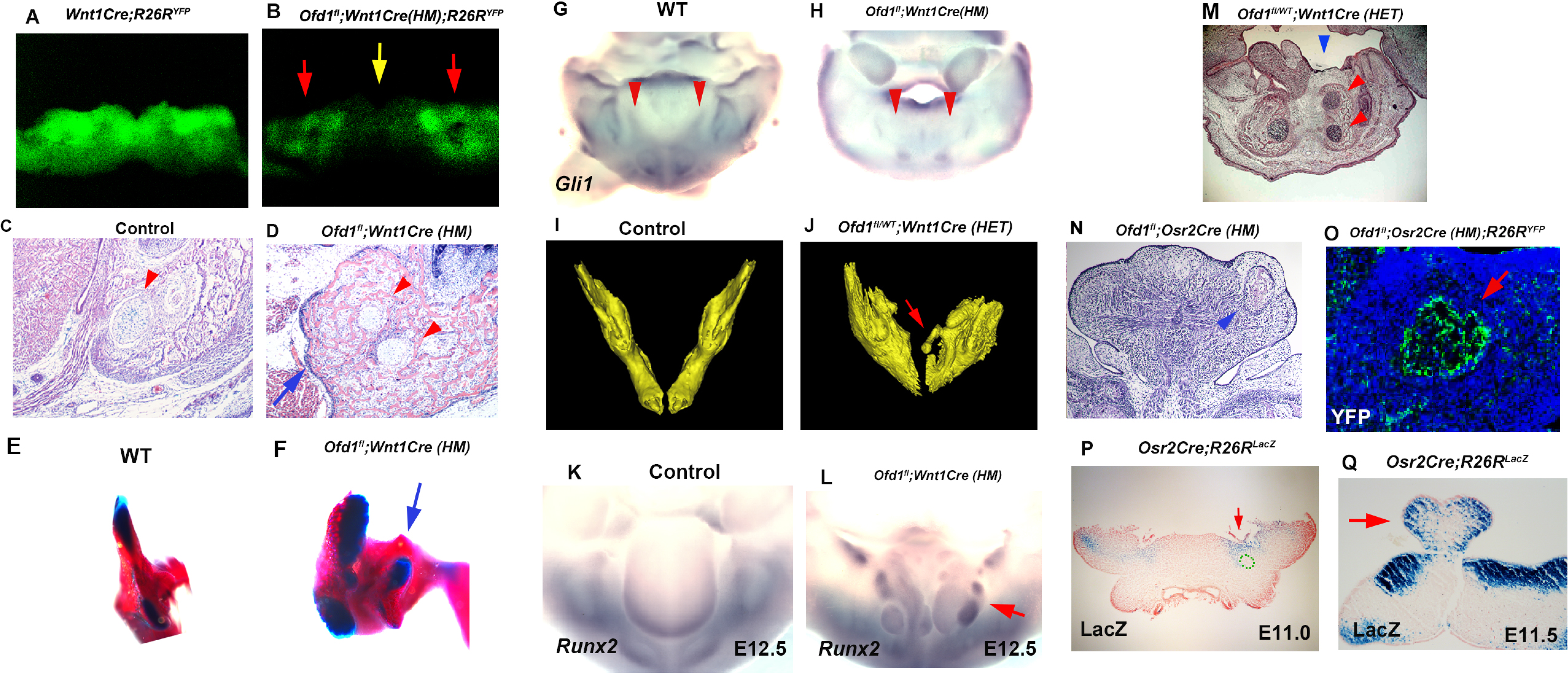
Bone in *Ofd1* mutant tongue. (A, B) Frontal images of YFP expression in *Wnt1Cre;R26R^YFP^* (A) and *Ofd1^fl^;Wnt1Cre(HM);R26R^YFP^* (B). Red and yellow arrows indicating developing mandibular bone region and early tongue swelling region, respectively. (C, D) Frontal section showing histological images in wild-type (C) and *Ofd1^fl^;Wnt1Cre(HM)* (D). Arrowhead indicating Meckele’s cartilage (C). Arrow and arrowheads indicating ectopic lingual bone and Meckel’s cartilage, respectively (D). (E, F) Posterior view of skeletal preparation of mandible in wild-type (E) and *Ofd1^fl^;Wnt1Cre(HM)* (F). Arrow indicating extra mandible. (G, H) Oral view of mandibles of whole mount *in situ* hybridization of *Gli1* in wild-type (G) and *Ofd1^fl^;Wnt1Cre(HM)* (H). Arrowheads indicating region correspond to ectopic lingual bone. (I, J) 3D reconstruction of mandibular bone of wild-type (I) and *Ofd1^fl/WT^;Wnt1Cre(HET)* (J) mice. Arrow indicating bone in the tongue (J). (K, L) Oral view of mandibles showing *in situ* hybridization of *Runx2* in wild-type (K) and *Ofd1^fl/WT^;Wnt1Cre(HET)* (L). Arrow indicating *Runx2* expression in the tongue (L). (M-Q) Frontal section showing histological images (M, N), immunohistochemistry of YFP (O) and LacZ staining (P, Q) in *Ofd1^fl/WT^;Wnt1Cre(HET)* (M), *Ofd1^fl/fl^;Osr2Cre(HM)* (N), *Ofd1^fl/fl^;Osr2Cre(HM);R26R^Yfp^* (O) and *Osr2Cre;R26R^LacZ^* (P, Q). Arrow indicating YFP expression in ectopic bone in the tongue (O). Arrows indicating LacZ positive region in lingual region of mandible (P) and tongue swelling (Q). Meckel’s cartilage was outlined by green dots (P).

### Cell migration for tongue frenum formation

The tongue consists of a main body and tongue frenum. The tongue frenum is often absent in *Ofd1^fl/WT^;Wnt1Cre(HET)* mice (see Fig. 1H). The tongue frenum is a thin flap of mucous membrane, and the tissue underneath the epithelium consist of CNCC, but not myogenic cells (Fig. 8A, 8B). The tongue frenum is already formed in wild-type mice at E12 when the tongue swelling becomes obvious (Fig. S9A), suggesting that the tongue frenum likely forms during emergence of the tongue swelling. It has been shown that tongue frenum is absent in *Lgr5* and *Tbx22* mutant mice (Morita et al., 2004, Pauws et al., 2009). We found restricted expression pattern of *Lgr5* and *Tbx22* in wild-type mandibular processes at E11.5 (Fig. 8C, 8D). *Lgr5* expression could not be detected in the mandibular processes at E10.5 (Fig. 8E, S9B). This indicated that the region showing *Lgr5* and *Tbx22* at E11.5 are probably involved in tongue frenum formation. Although the tongue frenum is located at the midline, *Tbx22* showed no its expression in the midline of the mandibular processes at E11.5 (Fig. 8D). The tongue frenum formation is unlikely to be relied on midline formation. In order to determine whether the anterior region of the *Lgr5*/*Tbx22*-expression domain is linked to tongue frenum formation, fate mapping analysis using DiD was performed in the wild-type mandible. DiD was injected into the anterior part of *Lgr5*/*Tbx22*-expressing region (AP region, Fig. 8F, 8G, S9C, S9D). DiD in the AP region moved to the midline region (Fig. 8G’, S9D’, S9D’’). Thus, cells at the AP region moved along the lingual-buccal axis and were then retained at the midline. When we injected DiO to more lateral regions of the *Lgr5*/*Tbx22*-expressing domain (LT region, Fig. S9C, S9D), cells labeled with DiO in the LT region moved along the anterior-posterior axis (Fig. S9D’, S9D’’). We already concluded that the TE region (the more posterior part of the mandibular processes from the AP regions at the midline) moved along the anterior-posterior axis (see Fig. 4Q-4Q’’). We then injected DiO at the TE region, and DiD at both left and right AP regions (Fig. 8H, 8I). Cells labeled with DiO (TE region) moved along the anterior-posterior axis and overtook cells with DiD (AP region) which moved across the lingual-buccal axis and merged at the midline (Fig. 8I’). Thus, distinct axis of cell migrations are present to form tongue swelling and frenum (Fig. 8J). To confirm whether cells in AP region form tongue frenum, we made incisions a lingual-buccal axis between the midline and AP region in wild-type mandibular processes to disturb cell migration from the AP region to the midline, and cultured embryos (Fig. 8K, 8K’). No tongue frenum was observed in cultured mandibles with incision (n=1/6, Fig. 8M), while the tongue frenum was present in most cultured mandibles without incision (n=8/10, Fig. 8L). Thus, cells in the AP region move the lingual-buccal axis to form the tongue frenum. Indeed, serial histological observation indicated that in some *Ofd1^fl/WT^;Wnt1Cre(HET)* mice, the tongue frenum was absent at the most anterior part of mandible (Fig. 8N), whilst the frenum was present at a slightly more posterior region in the same mouse (Fig. 8O; 8N and 8O were obtained from the same mutant mouse).

**Figure 8.**
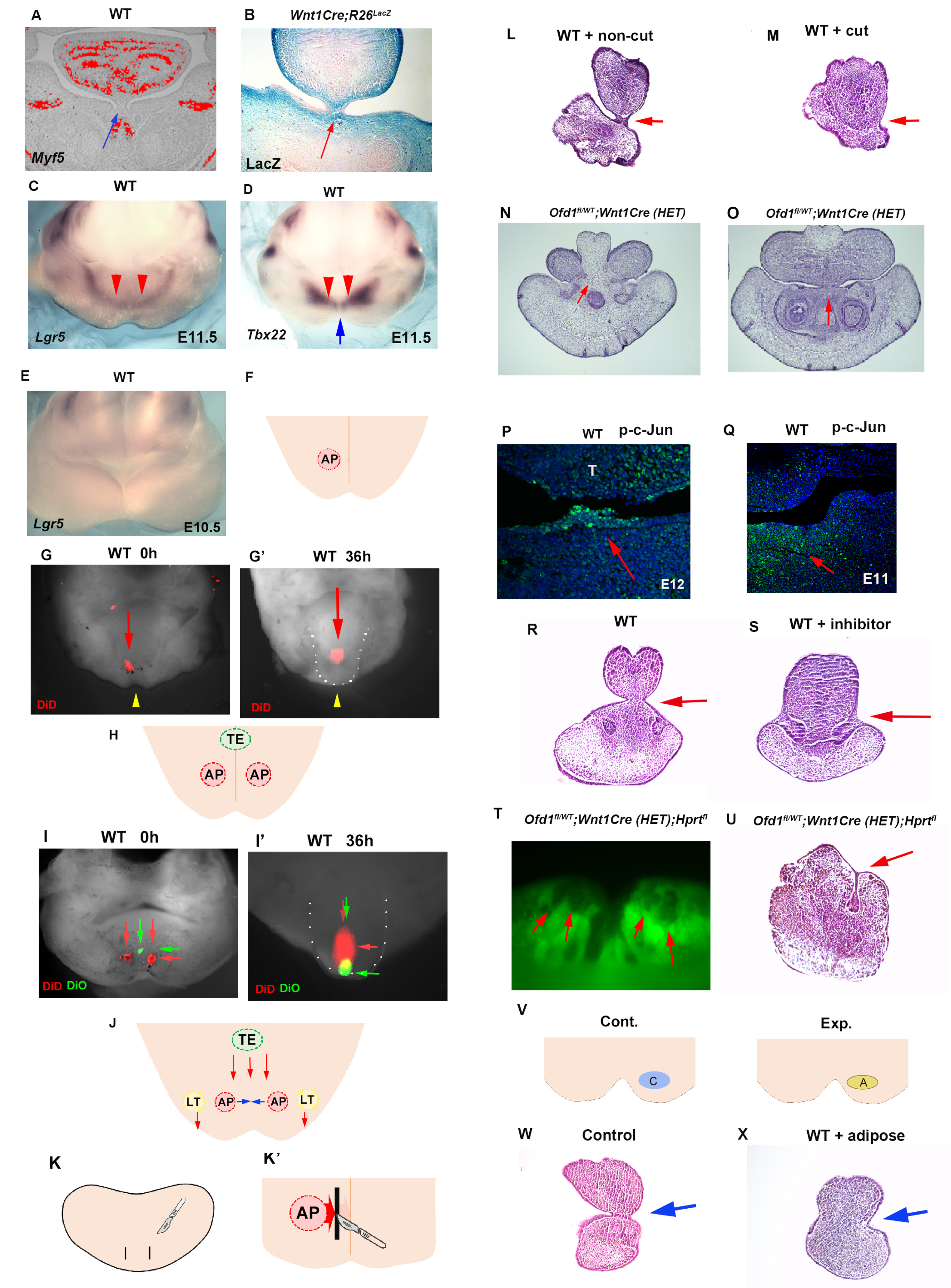
Tongue frenum in *Ofd1* mutant mice. (A, B) Frontal sections showing *in situ* hybridization of *Myf5* (A) and LacZ staining (B) in wild-type (A) and *WntCre;R26R^LacZ^* (B). Arrows indicating tongue frenum region. (C-E) Oral view of whole mount *in situ* hybridization of *Lgr5* (C, E) and *Tbx22* (D) in wild-type at E11.5 (C, D) and E10.5 (E). Red and blue arrows indicating anterior expression domain and midline region, respectively. (F, H) Scheme diagram showing oral view of mandible with AP (light red) and TE region (light gree), and midline (orange line). (G, G’, I, I’) Nuclear fluorescent image showing oral view of wild-type mandible before (G, I) and after (G’, I’) culture. Red and green arrows indicating DiD and DiO, respectively. Yellow arrowheads indicating midline. (G’, I’) Tongue like structure was partially removed and presumptive tongue-like structure are outlined by white dots. (J) Scheme diagram showing oral view of mandible with movement of cells in AP (light red), LT (light yellow) and TE region (light green line). (K, K’) Scheme diagram showing oral view of mandible with incision (black line) and midline (orange line). (L, M) Frontal sections showing histological images of cultured jaw without incision (N) and with incision (O). Arrows indicating the tongue frenum region. (N, O) Frontal sections showing histological images in *Ofd1^fl/WT^;Wnt1Cre(HET)* mice obtained from same mouse (O; more posterior site than N). (P, Q) Frontal sections showing immunohistochemistry of p-c-Jun in wild-type at E12 (P) and E11 (Q). Arrows indicating positive cells in the tongue frenum (P) and AP region (Q). (R, S) Frontal sections showing histological images of cultured jaw with (S) and without U73122 (R). Arrows indicating the tongue frenum region. (T) Oral view of mandible in *Ofd1^fl/WT^;Wnt1Cre(HET);Hprt^fl^* mice. Arrows indicating Gfp-negative domains around AP region before culture. (U) Frontal sections showing histological images of T after culture. Arrow indicating tongue frenum region. (V) Scheme diagram showing oral view of mandible with replacement (A; yellow circle=adipose, blue circle=CNCC). (W, X) Frontal sections showing histological images of cultured jaw without replacement (W) and with replacement (X). Arrows indicating tongue frenum region.

No activation of Hh or canonical Wnt signaling could be seen in the wild-type tongue frenum (Fig. S9E, S9F), suggesting that Hh or canonical Wnt signaling signaling are not involved in tongue frenum formation. In addition to *Lgr5*/*Tbx22*, it has been shown that the deletion of non-canonical Wnt signaling-related molecules, *Wnt5a* and *Ror2*, led to a lack of tongue frenum (Liu et al., 2012, Mossaad et al., 2018). We found p-c-Jun (marker of non-canonical Wnt signaling) positive cells in wild-type tongue frenum at E12, and in AP region at E11.5 (Fig. 8P, 8Q). Moreover, no tongue frenum formation was observed in wild-type mice, when the mandible was cultured with inhibitor of non-canonical Wnt signaling, U73122 (n=4/4; Fig. 8S). Thus, cell migration for tongue frenum formation are under control of non-canonical Wnt signaling. The presence or absence of the tongue frenum was varied between *Ofd1^fl/WT^;Wnt1Cre(HET)* mice, which is likely relied on X-inactivation. In fact, when we cultured mandibles of *Ofd1^fl/WT^;Wnt1Cre(HET);Hprt^GFP^* mice showing cells with inactivation of normal X-chromoseome (Gfp-negative) at the AP region (Fig. 8T; before culture), the tongue frenum was absent (Fig. 8U; after culture). The Gfp-negative domain in *Ofd1^fl/WT^;Wnt1Cre(HET);Hprt^GFP^* mice were found to contain adipocytes. To understand whether the presence of adipose at AP region affect tongue frenum formation, CNCC were replaced to adipose at AP region in wild-type mice, and cultured them. The tongue frenum was impaired in the cultured wild-type mandible with replacement (n=4/5; Fig. 8W, 8Y). Thus, tongue frenum formation is also reliant on which type of cells firstly reach to the AP region. Tongue frenum formation occurred when CNCC reach first, while it is inhibited when adipocytes reach first.

## Discussion

Tongue development contains multiple cellular processes including cell-cell interaction, migration and differentiation. We found that contact between CNCC and mesoderm-derived cells within hypoglossal cord are involved in differentiation of mesoderm-derived cells into myoblasts through Hh signaling in CNCC (Fig. 9). In addition, CNCC in lingual region of mandibular processes require Hh signaling activity for their own differentiation into fibroblasts to form tongue connective tissue. Hh signaling activity in CNCC is also essential for their migration to form tongue swelling, but not for their migration to form mandibular bone or tongue connective tissue. Moreover, tongue morphology is determined by which type of cells firstly reach to the specific region, that affect subsequent cell migration. There are multiple cell migration axis in mandibular processe to form tongue main body and tongue frenum. Multiple cellular processes are thus required to be coordinated in distinct region and timing during tongue development. These tongue anomalies are observed in syndromic and non-syndromic condition in human (Cobourne et al., 2019, Hill et al., 2020, Li et al., 2020, Yin et al., 2020). Our findings revealed that genetical or mechanical disturbances in these processes results in syndromic or non-syndromic tongue anomalies, respectively. Moreover, these cellular processes are under the control of X-inactivation. Thus, tongue development is an excellent model to investigate how multiple cellular processes are coordinated during organogenesis.

**Figure 9.**
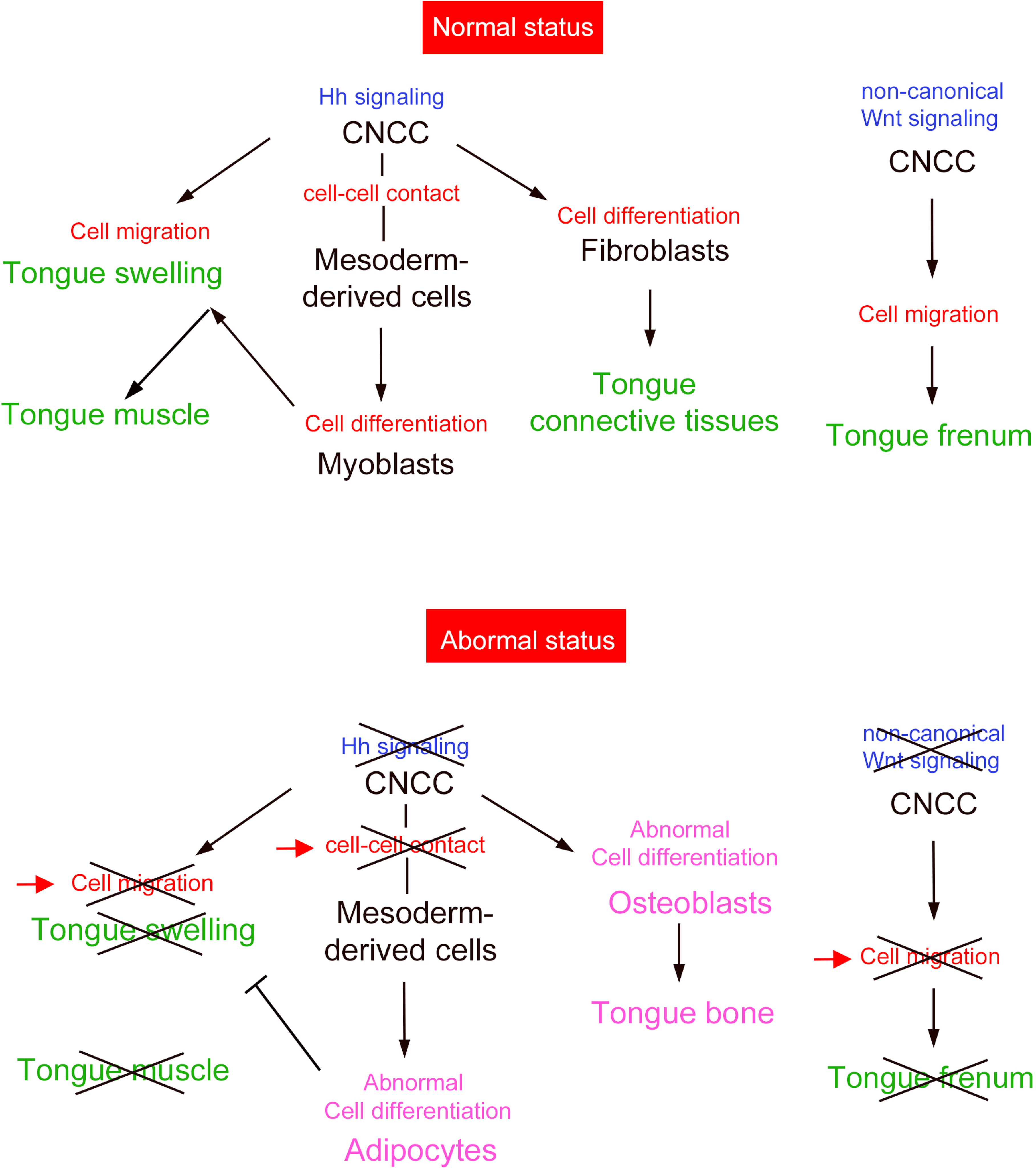
Summary. Tongue development require coordinated multiple cellular processes.

In *Ofd1^fl^;Wnt1Cre(HM)* mice, CNCC was absent in the region where tongue swelling emerged, while CNCC was present in mandibular bone and its surrounding region, although all CNCC have no *Ofd1*. There are two types of CNCC (for tongue swelling and tongue connective tissue).

The phenotypic variation seen in the tongues of *Ofd1^fl/WT^;Wnt1Cre(HET)* mice is likely caused by X-inactivation. However, *Ofd1^fl/WT^;Osr2Cre(HET)* mice display no apparent brown adipose tissue, even though X-inactivation is present in these mice. Instead, brown adipose tissue was observed in all *Ofd1^fl^;Osr2Cre(HM)* mice, in which no X-inactivation occurs. All cells in the *Osr2Cre* expression domain were *Ofd1* mutant cells in *Ofd1^fl^;Osr2Cre(HM)* mice, whereas only a certain number of cells were *Ofd1* mutant cells in *Ofd1^fl/WT^;Osr2Cre(HET)* mice, due to X-inactivation. The *Osr2Cre* domain only partially overlaps with the hypoglossal cord (see Fig. 2N’’). These results indicate that *Ofd1* deletion from all cells of the *Osr2Cre* expression domain is necessary for obvious brown adipose tissue formation, while *Ofd1* deletion from some cells within the *Osr2Cre* expression domain is not sufficient to induce morphological detectable brown adipose tissue formation. *Ofd1* mutation always occurred in the same region within the hypoglossal cord in *Ofd1^fl^;Osr2Cre(HM)* mice, as X-inactivation did not occur in these mice. However, the location of adipose tissue in the tongues of *Ofd1^fl^;Osr2Cre(HM)* mice varied between individuals. It has been shown that CNCCs act as a scaffolding structure for the organization of migrating myoblasts. Furthermore, the direction of cell migration is determined after CNCCs and myoprogenitors reach to the mandible. Thus, the presence of cells with *Ofd1* mutation disorganize migration of CNCC, myogenic progenitors and ectopic adipocytes.

Ciliopathies show prominent mixed symptoms in several organs including the tongue. Our findings provide hints for possible future treatment in ciliopathies.

## Materials and Methods

### Production and analysis of transgenic mice

All animal experiments were reviewed and approved by the Niigata University Institutional Animal Care and Use Committee (approval number SA00610, SD01308). *Ofd1^fl/fl^*, *Smo^fl/fl^*, *R26R^LacZ^*, *R26R^Yfp^*, *Wnt1Cre*, *Osr2Cre*, *K14Cre* and *Ift88^fl/fl^* mice were produced as described by Ferrante et al., (2006), Jeong et al., (2004), Soriano (1999), Srinivas et al., (2001), Danielian et al. (1998), Lan et al., (2007), Yi et al., (2006) and Haycraft et al., (2007), respectively. *Hprt^fl^* mice were purchased from the Jackson laboratory. Embryonic day 0 (E0) was taken to be midnight prior to finding a vaginal plug.

### *In situ* hybridization

*In situ* hybridization was carried out to detect mRNAs using [^35^S]UTP or DIG as described previously (Ohazama et al., 2008).

### Immunohistochemistry

Frozen sections were incubated with antibodies to GFP (ab13970; Abcam), p-c-Jun (#9261; Cell signal), KRT1-5 (14309-1-AP ; Proteintech), Prdm16 (AF6295; R&D), UCP1 (U6382; Sigma) and MyoD (M351201; Dako). To detect anti-GFP or anti-MyoD antibodies, the sections were incubated with Alexa488-conjugated secondary antibody. A tyramide signal amplification system (Perkin Elmer Life Science, Waltham, MA, US) was used to detect the anti-KRT1-5 and anti-p-c-Jun antibodies.

### Quantitative-PCR (Q-PCR)

E16.5 embryos were frozen, and were sectioned into 12μm thick slices. Then, sections were mounted on PEN membrane slides, which were stained by Toluidine blue. Sparse region were dissected with the Laser micro dissection system (Leica Microsystems) into a microcentrifuge tube cap placed directly beneath the section. The tube cap was filled with 75 μl of RNAlater (Sigma-Aldrich). RNA was isolated using a RNeasy Mini Kit (Qiagen). Q-PCR was performed using GoTaq qPCR Master Mix (Promega) with the carboxy-X-rhodamine (CXR) Dye and Rotor-Gen-Q (Qiagen) detection system. All samples were run in triplicate for each experiment, and relative transcript abundance was normalized to the amount of GAPDH.

### 3D reconstruction of mandibular bone

Heads of *Ofd1^fl/WT^;Wnt1Cre(HET)* and wild-type mice were scanned with an Explore Locus SP (GE Pre-clinical Imaging) high-resolution micro-CT with a voxel dimension of 8 µm. 3D reconstruction was performed using Microview (GE Pre-clinical Imaging) and Mimics (Materialise).

### Human sample

Approval for study on human subjects was obtained from the Niigata University (2018-0228). Informed consent was obtained to use tongue tissue. Clinical information including a three-generation pedigree was obtained. DNA was extracted using blood sample. Mutations in OFD1 were screened by using primer sequences covering all 23 exons. Two surgical specimens were selected from the surgical pathology files in the Division of Oral Pathology, Niigata University Graduate School of Medical and Dental Sciences.

### Organ culture

The mandible including the tongue region was separated from the head and body at E11.5. The mandible explants were immediately placed into a glass bottle containing culture media comprising 100% immediately centrifuged (IC) rat serum. IC rat serum was incubated at 56°C for 30 min to inactivate complement. The culture bottles were attached to a rotator drum and rotated at 14 rpm at 37°C while being continuously supplied with 5% O2/ 95% CO2 gas mixture. At the end of the culture period, explants were fixed with 4% PFA, and processed for histological examination. U73122 (Selleck Chemicals) was used as 10µM.

### Culture of human white adipocyte and myoblasts

Human skeletal muscle myoblasts (CC-2561) and subcutaneous preadipocyte cells (PT-5020) were purchased from Lonza (Lonza Walkersville). Subcutaneous preadipocyte cells were induced to differentiate into terminal white adipocytes according to the manufacturer’s protocol.

### Pseudo scanning electron microscopy (SEM)

Pseudo SEM analysis was carried out as described previously (Sandell et al., 2012). Briefly, Separated maxilla and mandible were fixed with 4% PFA, and then, stained with DAPI. Images were taken by confocal microscopy. The multiple sections were merged into a single projection view to generate the pseudo SEM image.

### Statistical analysis

Excel Toukei (ver. 6.0) was used for statistical analysis, which was done with a two-tailed unpaired Student’s t test. P<0.05 was considered statistically significant.

## Acknowledgments

This research was funded by the Japan Society for the Promotion of Science (JSPS; 17H01601, 20K18753, 20K18661, 21K10088).

## Author contributions

MK, KK, FTS, TK, JN, MK, TN, VU, YI, FM, TN, YK, SM, J-IT, YK, TM, RHK, PC, PTS, MTC, BF performed experiments and analysed the data. AO design experiments, interpreted the data and wrote the manuscript. All authors reviewed and edited the manuscript.

---

**Supplementary Figure 1.**
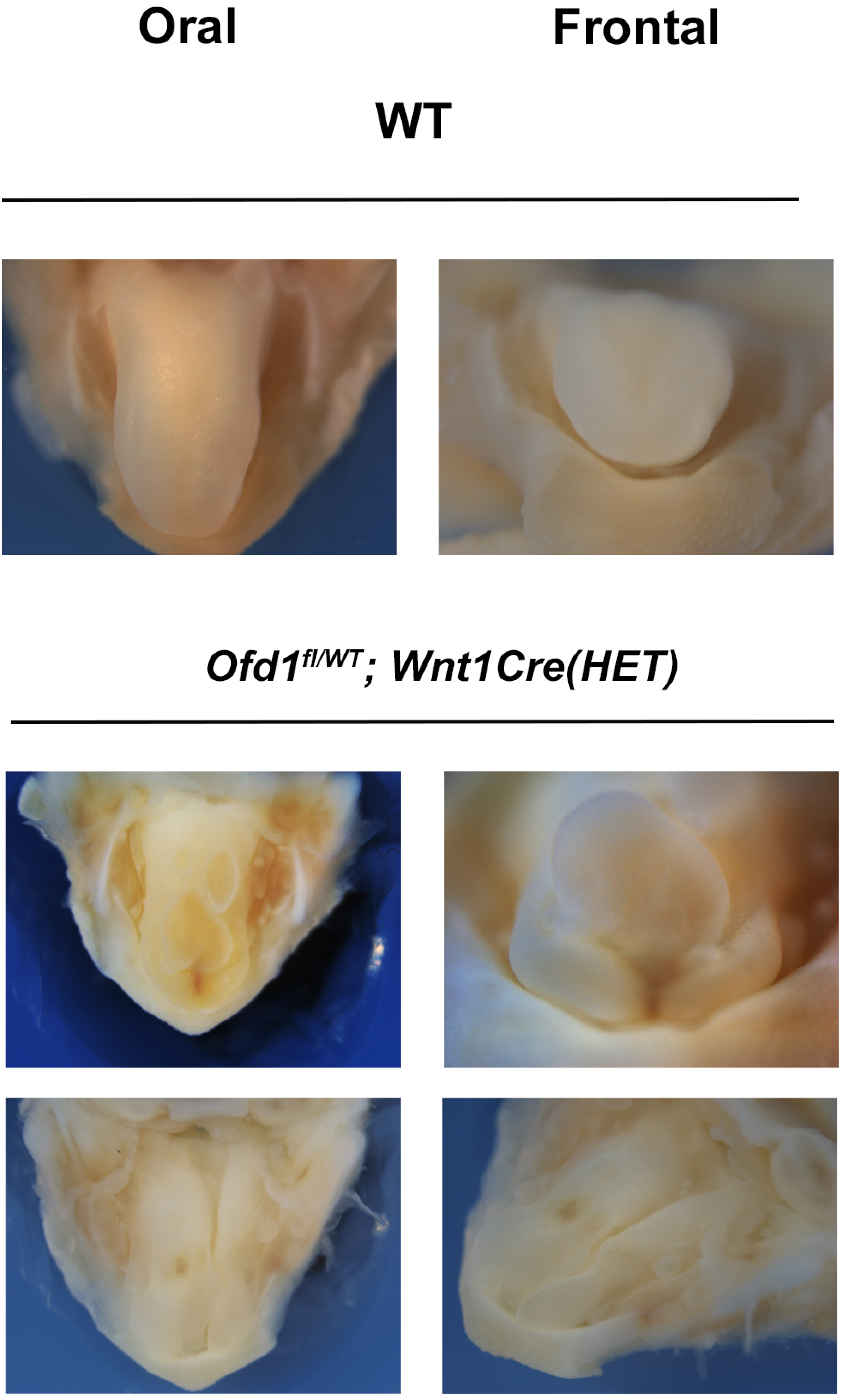
Abnormal shaped tongue in *Ofd1^fl/WT^;Wnt1Cre(HET)* mice. Oral and frontal view of the abnormal shaped tongue in *Ofd1^fl/WT^;Wnt1Cre(HET)* mice

**Supplementary Figure 2.**
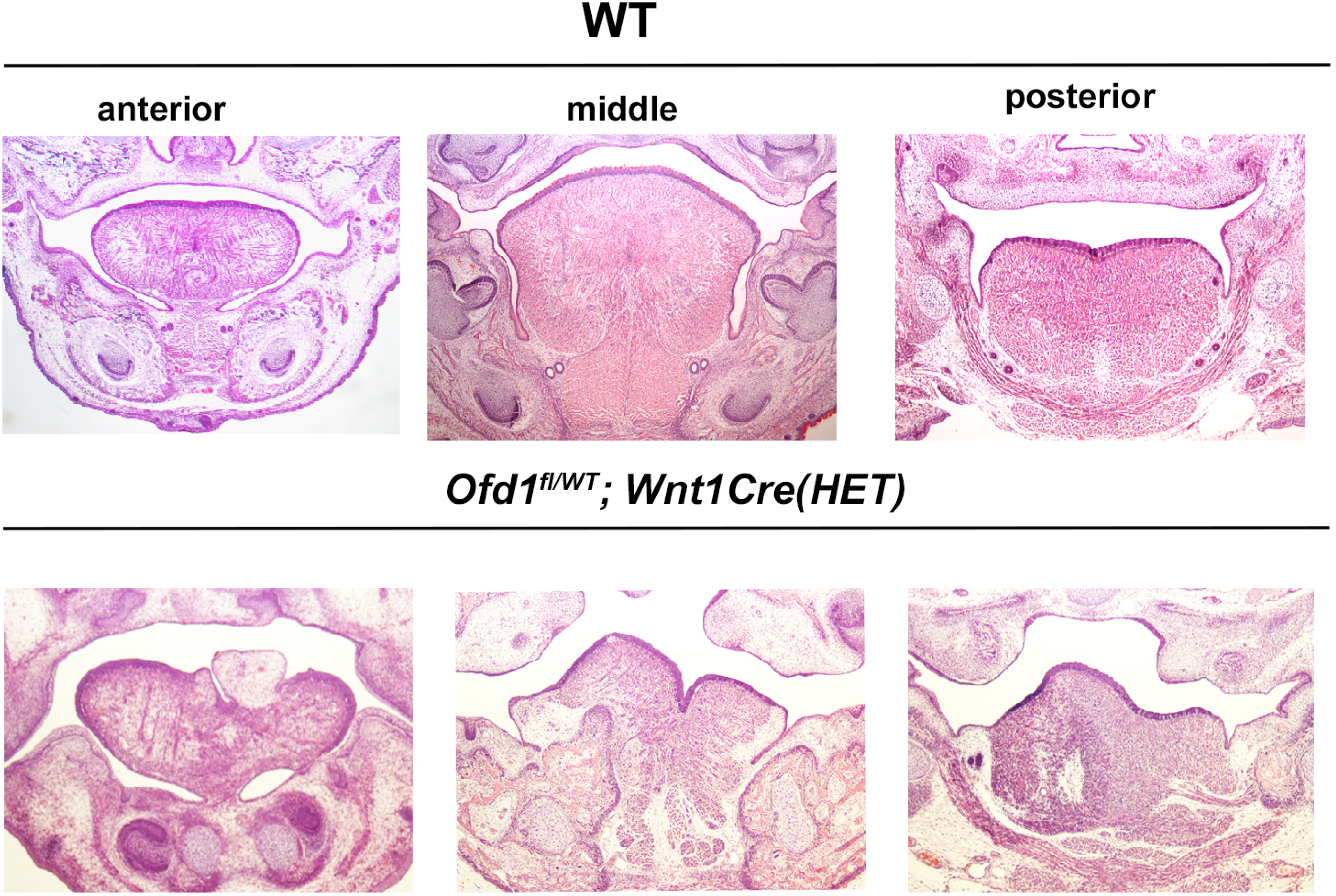
Tongue phenotypes in *Ofd1^fl/WT^;Wnt1Cre(HET)* mice. Frontal section showing histological images of tongue in wild-type and *Ofd1^fl/WT^;Wnt1Cre(HET)* mice at E18.5.

**Supplementary Figure 3.**
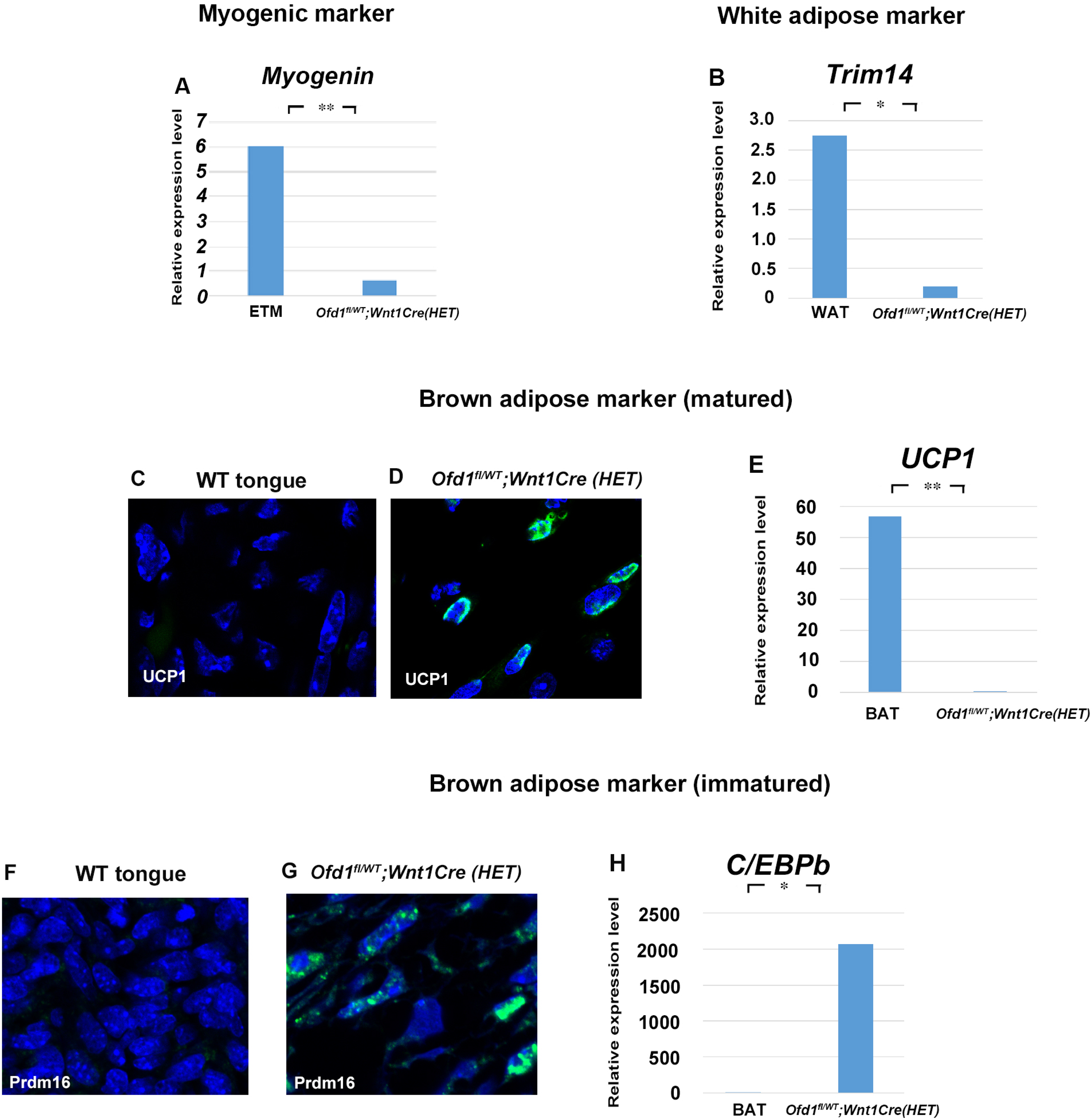
qPCR and immunohistochemistry analysis on *Ofd1^fl/WT^;Wnt1Cre(HET)* mice. (A, B) q-PCR on mRNA isolated from ectopic sparse tissue from *Ofd1^fl/WT^;Wnt1Cre(HET)*, embryonic tongue muscle (ETM) and white adipose tissue (WAT). ** P<0.01. Only low-level expression of myogenic markers in the sparse tissue compared with the embryonic tongue muscle (A). Expression levels of white adipocyte markers in the sparse tissue were significantly lower than those in white adipose, adult inguinal fat (B). (C, D) Frontal sections showing immunohistochemistry of marker of matured brown adipose tissue (UCP1). UCP1 was expressed in the sparse tissue of *Ofd1^fl/WT^;Wnt1Cre(HET)* mice, whereas no expression was observed in wild-type tongue. (E) q-PCR on mRNA isolated from ectopic sparse tissue from *Ofd1^fl/WT^;Wnt1Cre(HET)* and brown adipose tissue (BAT). Expression of UCP1 in the sparse tissue was lower than that in perirenal fat (Perirenal fat is mainly composed of matured brown adipocytes). (F, G) Frontal sections showing immunohistochemistry of marker of immatured brown adipose tissue (Prdm16). Prdm16 was expressed in the sparse tissue of *Ofd1^fl/WT^;Wnt1Cre(HET)* mice, whereas no expression was observed in wild-type tongue. (H) q-PCR on mRNA isolated from ectopic sparse tissue from *Ofd1^fl/WT^;Wnt1Cre(HET)* and matured brown adipose tissue (BAT; perirenal fat). Expression of another marker of immatured brown adipose tissue, *C/EBPb*, in the sparse tissue was higher than that in matured brown adipocytes, indicating that the sparse tissue contained many pre-brown adipocytes in comparison with matured brown adipose tissue.

**Supplementary Figure 4.**
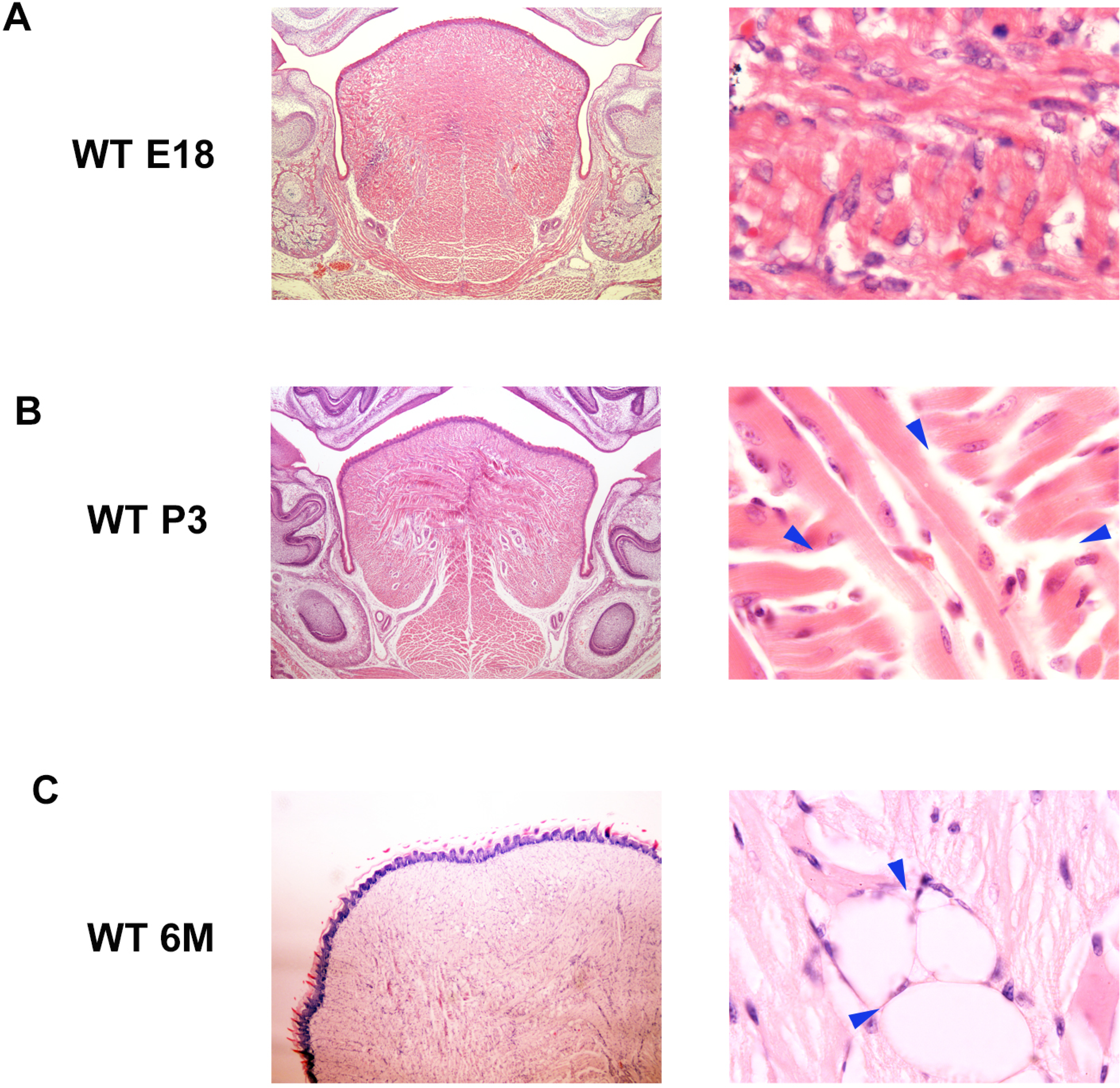
Artifact and adipose tissue in wild-type tongue. (A, B) Frontal sections showing empty spaces without surrounding cells in the tongue of prenatal (A) and postnatal (B) wild-type mice. Right panels are high magnification of left panels. Arrowheads indicating empty spaces. (C) Frontal sections showing intermuscular adipose in the tongue of 6-month-old wild-type mice. Arrowheads indicating adipocytes. P3; postnatal day 3, 6M; 6-month-old. Mucous salivary glands and their associated-adipose are known to be present in wild-type tongue. However, we could not identify any epithelial cells including glandular bodies or excretory ducts through immunohistochemistry around the sparse tissue. Adipose is also known to be present in the tongue as intermuscular adipose, which is often pathologically enlarged (Colella et al., 2009). However, it is known that the intermuscular adipose in the tongue is related to aging (Vettor et al., 2009, Scarda et al., 2010). In prenatal or postnatal wild-type tongue, empty gaps were found between muscle bundles; however, adipocyte-like cells could not be observed around the gaps (Fig. S4A, S4B). However, empty gaps surrounded by cells with a thin rim of cytoplasm between muscle bundles were observed in older mice (Fig. S4C). Empty gaps between neighboring muscle fibers found in prenatal or postnatal wild-type tongue should be an artifact, as previously reported (Rother et al., 2002). Thus, the brown adipose tissue found in the *Ofd1* mutant tongue was not related to mucous salivary glands associated-adipose or intermuscular adipose.

**Supplementary Figure 5.**
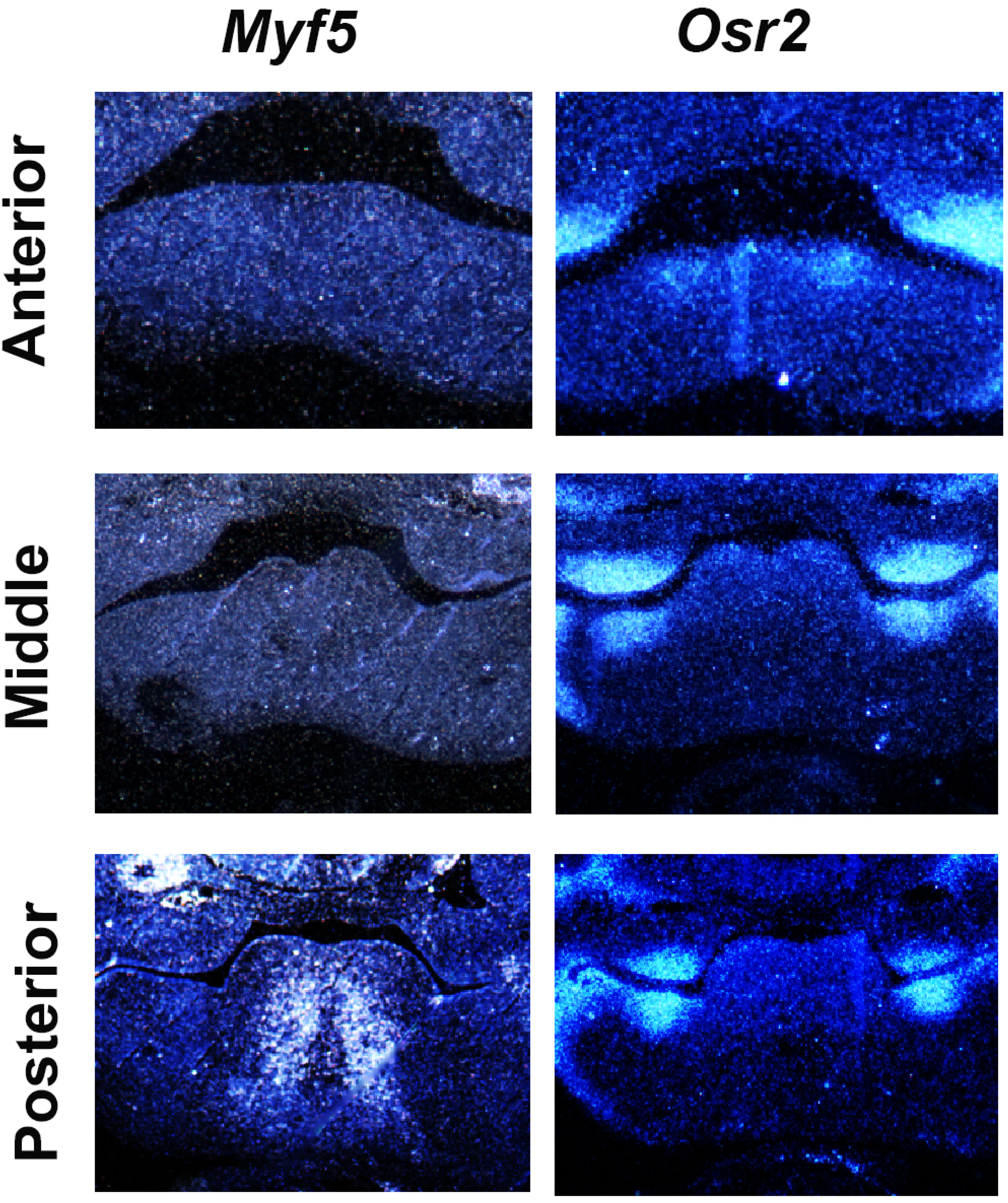
*Myf5* and *Osr2* expression in developing tongue. Frontal sections showing *in situ* hybridization of *Myf5* and *Osr2* in wild-type mice at E11.5.

**Supplementary Figure 6.**
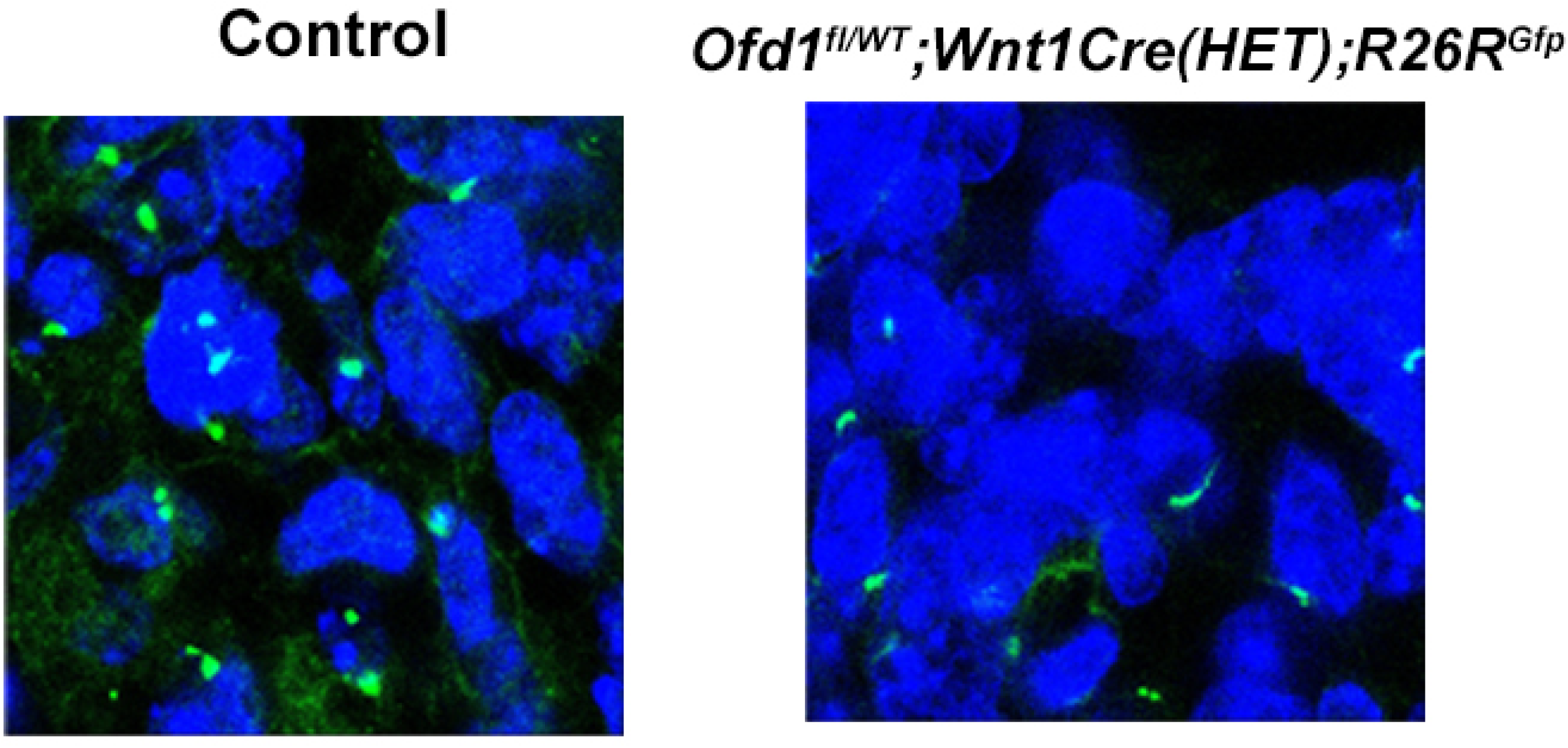
Acetylated α-tubulin in CNCC during tongue development. Frontal section showing immunohistochemistry of acetylated α-tubulin in YFP-expressing cells of *Wnt1Cre(HET);R26R^Yfp^* and *Ofd1^fl/WT^;Wnt1Cre(HET);R26R^Yfp^* mice. CNCC were confirmed by YFP expression.

**Supplementary Figure 7.**
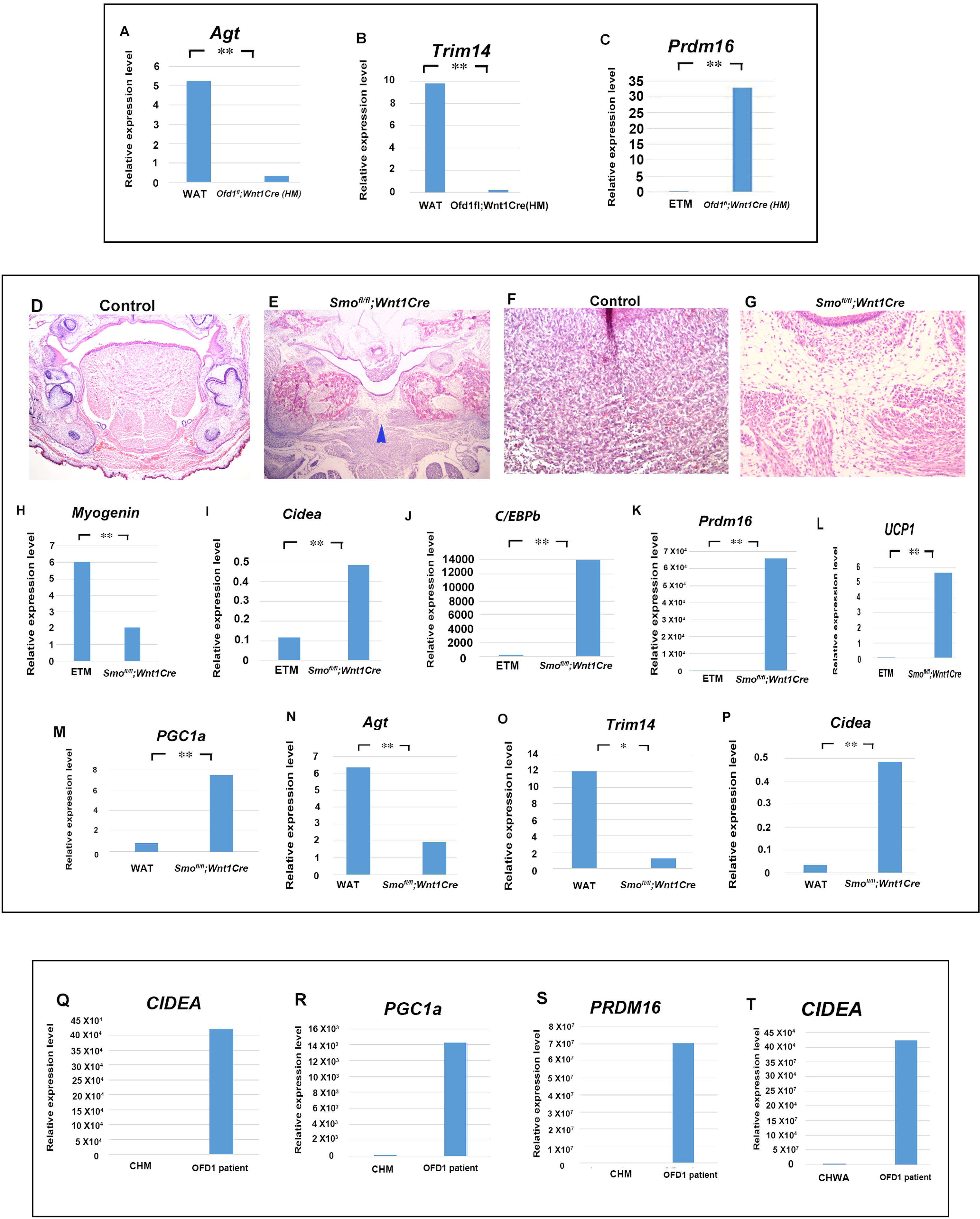
qPCR analysis on ectopic sparse tissue. (A-C) q-PCR on mRNA isolated from ectopic sparse tissue from *Ofd1^fl^;Wnt1Cre(HM)*, embryonic tongue muscle (ETM) and white adipose tissue (WAT). (D-G) Frontal section showing histological images of wild-type (D, F) and *Smo^fl/fl^;Osr2Cre(HM)* (E, G) mice. F and G were high magnification of D and E, respectively. (H-P) q-PCR on mRNA isolated from ectopic sparse tissue from *Smo^fl/fl^;Wnt1Cre*, embryonic tongue muscle (ETM) and white adipose tissue (WAT). (Q-T) q-PCR on mRNA isolated from ectopic sparse tissue from OFD1 patient tongue, cultured human myoblasts (CHM) and cultured human white adipocytes (CHWA).

**Supplementary Figure 8.**
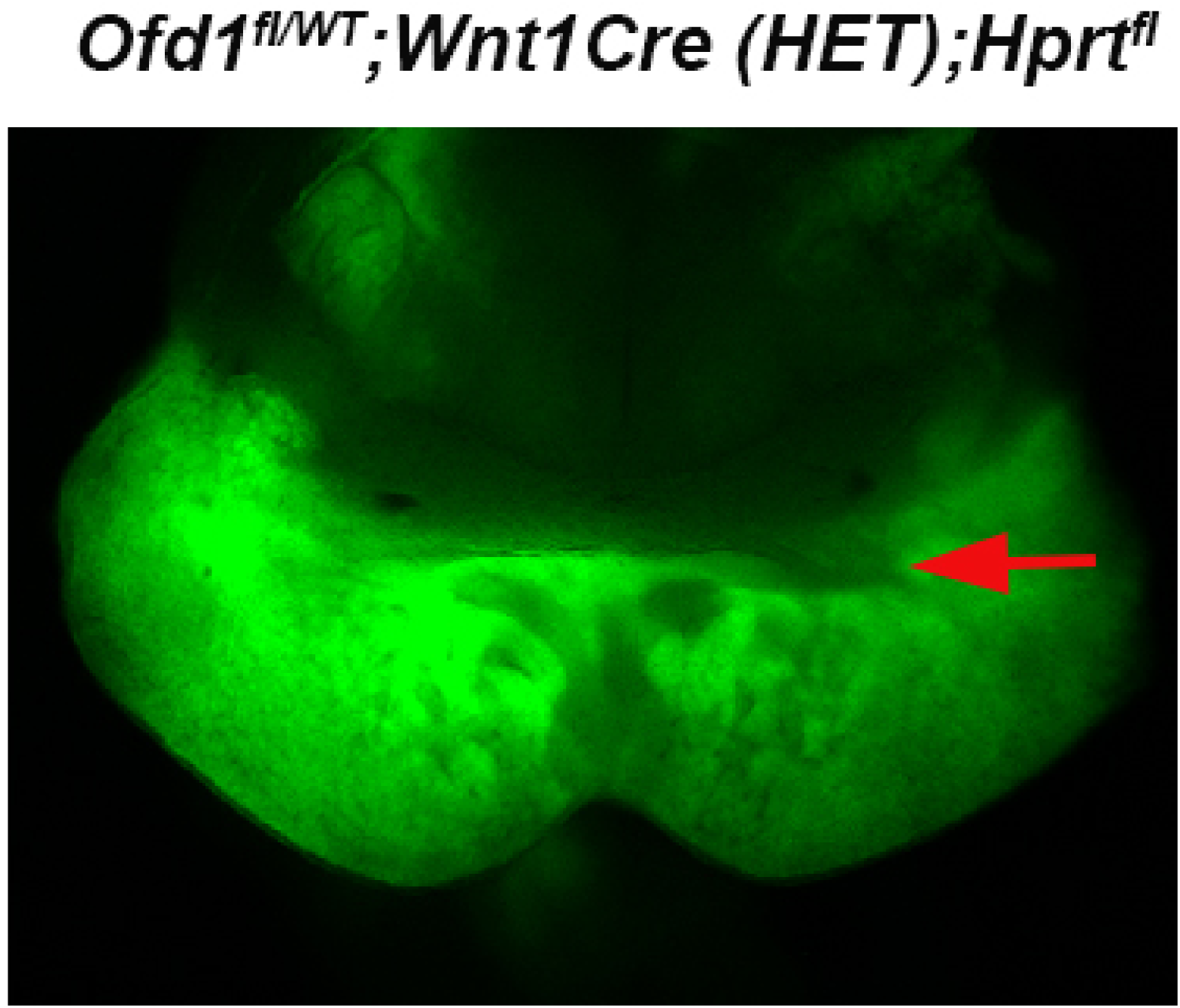
Osr2Cre in developing tongue. Oral view of mandible in *Ofd1^fl/WT^;Wnt1Cre(HET);Hprt^fl^* at E11.5. Arrow indicating Gfp-negative domain.

**Supplementary Figure 9.**
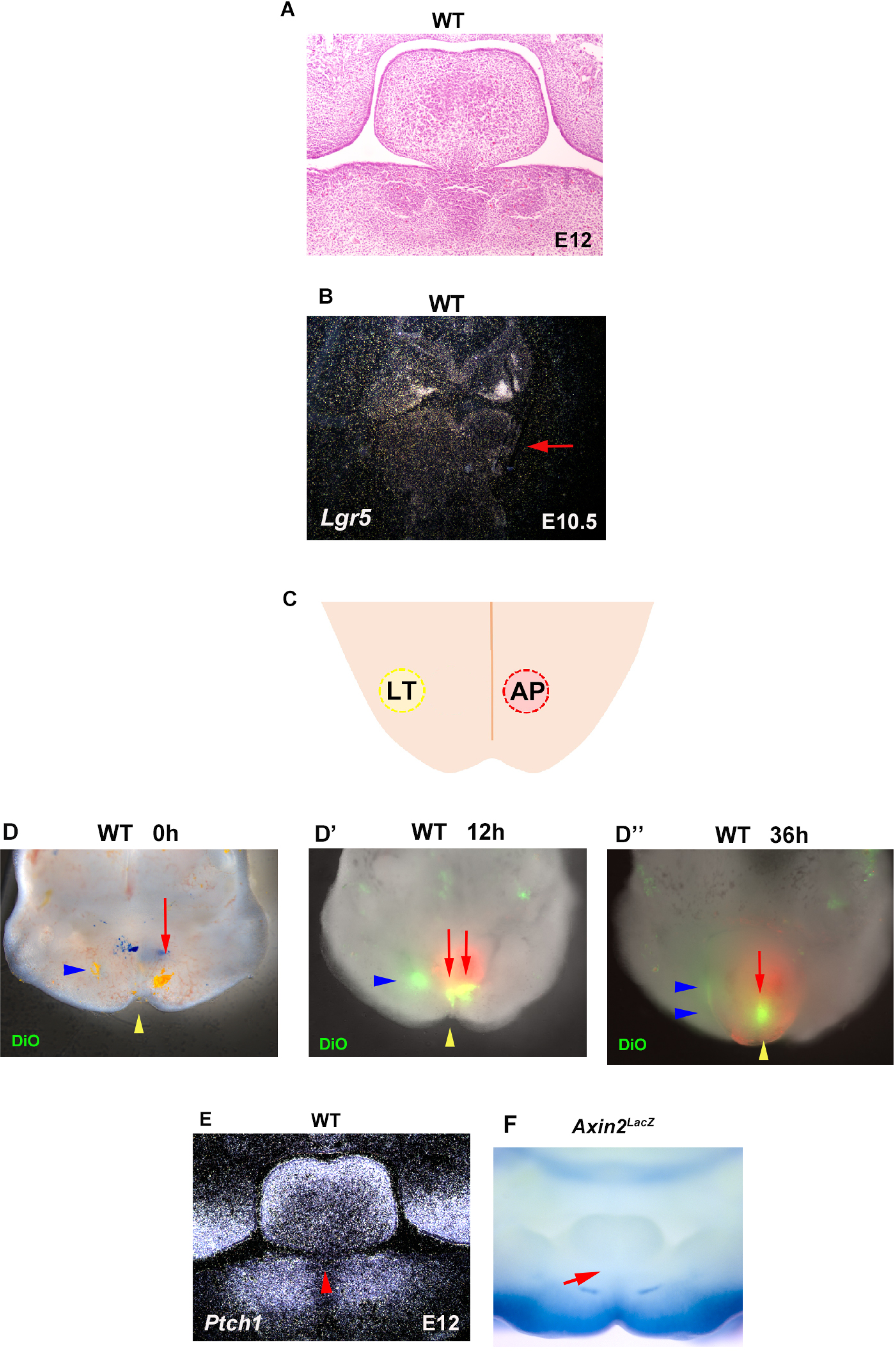
Tongue frenum formation in wild-type mice. (A, B) Frontal section showing histological image (A) and *in situ* hybridization of *Lgr5* (B) of tongue frenum (A) and mandible (B) in wild-type mice at E10.5 (B) and E12.5 (A). Arrow indicating mandible region. (C) Scheme diagram showing oral view of mandible with LT (yellow ceircle) and AP (right red circle) region, and midline (orange line). (D-D’’) Oral view of wild-type mandible with DiO injection (E; 0 h, E’; 12 h, E’’; 36 h after injection). Blue arrowheads and red arrows indicating DiO at LT and AP region, respectively. Yellow arrowhead indicating midline. (E) Frontal sections showing *in situ* hybridization of *Ptch1* at E12.5. (F) Frontal view of mandible in LacZ stained *Axin2^LacZ^* mice at E12.5. Arrows and arrowhead indicating tongue frenum region (E, F).

**Supplementary information 1.**
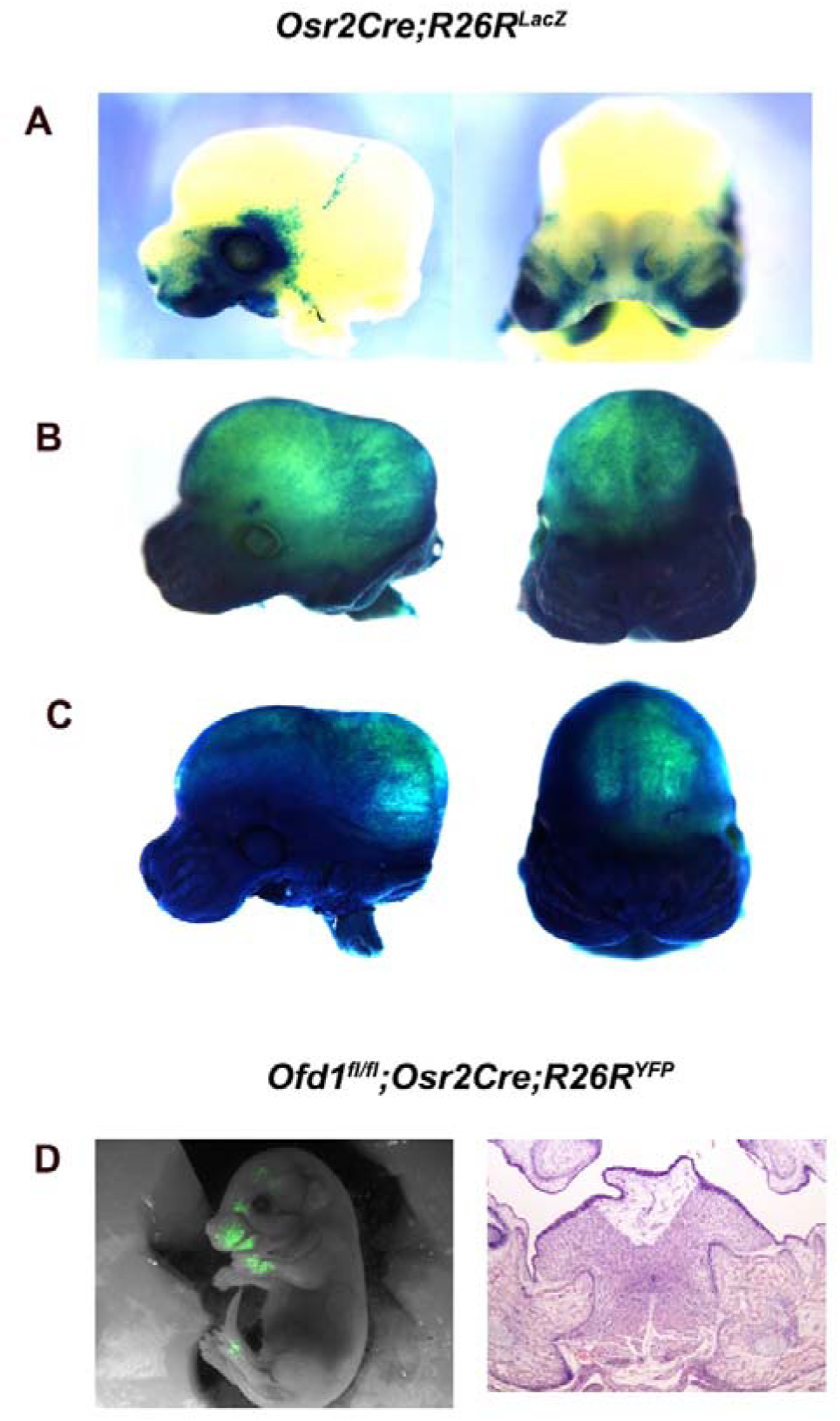
*Osr2Cre* activation (confirmed by LacZ staining in *Osr2Cre;R26R^LacZ^*) exhibited the three different patterns (Lan et al., 2007). Most embryos displayed tissue-specific pattern consistent with specific Cre expression from the *Osr2* locus, as shown in A (tissue-specific pattern). Approximately 20% of *Osr2Cre* displayed some ectopic activation, as shown in B (ectopic pattern). Approximately 10% of *Osr2Cre* embryos displayed almost ubiquitous activation, as shown in C (ubiquitous pattern). Abnormal tongue was observed in *Ofd1^fl^;Osr2Cre;R26R^YFP^* mice with tissue-specific pattern (D).

## References

1. Cobourne M, et al. How to make a tongue: Cellular and molecular regulation of muscle and connective tissue formation during mammalian tongue development. Semin Cell Dev Biol. 2019;91:45–54.

2. Hill RR, Lee CS, Pados BF. The prevalence of ankyloglossia in children aged <1 year: a systematic review and meta-analysis. Pediatr Res 2020);Nov,13.

3. Li J, et al. Giant sublingual hamartoma with medial cleft tongue: a case report and literature review. J Int Med Res 2020;48,300060520942089.

4. Yin S, Zhao Z. Incomplete Cleft Palate, Bifid Tongue, and Oral Hamartomas: A Rare Congenital Anomaly. J Craniofac Surg 2020;31,e184–e185.

5. Auber, B. et al. A disease causing deletion of 29 base pairs in intron 15 in the MKS1 gene is highly associated with the campomelic variant of the Meckel-Gruber syndrome. Clin Genet 72, 454–9 (2007).

6. Barbosa, A.C. et al. Hand transcription factors cooperatively regulate development of the distal midline mesenchyme. Dev Biol 310, 154–68 (2007).

7. Barron, F. et al. Downregulation of Dlx5 and Dlx6 expression by Hand2 is essential for initiation of tongue morphogenesis. Development 138, 2249–59 (2011).

8. Bhattacharya, V., Khanna, S., Bashir, S.A., Kumar, U. & Garbyal, R.S. Cleft palate associated with hamartomatous bifid tongue. Report of two cases. J Plast Reconstr Aesthet Surg 62, 1442–5 (2009).

9. Bisgrove, B.W. & Yost, H.J. The roles of cilia in developmental disorders and disease. Development 133, 4131–43 (2006).

10. Brugmann, S.A. et al. A primary cilia-dependent etiology for midline facial disorders. Hum Mol Genet 19, 1577–92 (2010).

11. Cobourne M, Sachiko Iseki c, Anahid A. Birjandia, Hadeel Adel Al-Lamia,d, Christel Thauvin-Robinete,f, Guilherme M. Xaviera,b, Karen J. Liua How to make a tongue: Cellular and molecular regulation of muscle and connective tissue formation during mammalian tongue development. Semin Cell Dev Biol. 2019 Jul;91:45–54.

12. Colella, G. et al. Giant intramuscular lipoma of the tongue: a case report and literature review. Cases J 2, 7906 (2009).

13. Danielian, P.S., Muccino, D., Rowitch, D.H., Michael, S.K. & McMahon, A.P. Modification of gene activity in mouse embryos in utero by a tamoxifen-inducible form of Cre recombinase. Curr Biol 8, 1323–6 (1998).

14. Daramola, O.O., Suchi, M. & Chun, R.H. Lingual hamartoma associated with a cleft palate in a newborn. Ear Nose Throat J. 93, E9–11 (2014).

15. Depew, M.J. et al. Dlx5 regulates regional development of the branchial arches and sensory capsules. Development 126, 3831–46 (1999).

16. do Rêgo Barros de Andrade Fraga, M., Azoubel Barreto, K., Barbosa Lira, T.C. & Aparecida de Menezes, V. Is the Occurrence of Ankyloglossia in Newborns Associated with Breastfeeding Difficulties? Breastfeed Med 15, 96–102 (2020).

17. Ferrante, M.I. et al. Identification of the gene for oral-facial-digital type I syndrome. Am J Hum Genet 68, 569–76 (2001).

18. Ferrante, M.I. et al. Convergent extension movements and ciliary function are mediated by ofd1, a zebrafish orthologue of the human oral-facial-digital type 1 syndrome gene. Hum Mol Genet 18, 289–303 (2009).

19. Ferrante, M.I. et al. Oral-facial-digital type I protein is required for primary cilia formation and left-right axis specification. Nat Genet 38, 112–7 (2006).

20. Goggolidou, P. Wnt and planar cell polarity signaling in cystic renal disease. Organogenesis 10, 86–95 (2014).

21. Gorini E, Mullace M, Migliorini L, Mevio E. Osseous choristoma of the tongue: a review of etiopathogenesis. Case Rep Otolaryngol. 2014;2014:373104.

22. Han, A., Zhao, H., Li, J., Pelikan, R. & Chai, Y. ALK5-mediated transforming growth factor β signaling in neural crest cells controls craniofacial muscle development via tissue-tissue interactions. Mol Cell Biol 34, 3120–31 (2014).

23. Haycraft, C.J. et al. Intraflagellar transport is essential for endochondral bone formation. Development 134, 307–16 (2007).

24. Hemmaoui, B., Sahli, M., Jahidi, A. & Benariba, F. Hamartoma of the tongue. Eur Ann Otorhinolaryngol Head Neck Dis 134, 295–296 (2017).

25. Hill, R.R., Lee, C.S. & Pados, B.F. The prevalence of ankyloglossia in children aged <1 year: a systematic review and meta-analysis. Pediatr Res (2020).

26. Hiradfar, M. et al. Accessory tongue: Classification and report of a case. Int J Pediatr Otorhinolaryngol 79, 1175–9 (2015).

27. Hosokawa, R. et al. TGF-beta mediated FGF10 signaling in cranial neural crest cells controls development of myogenic progenitor cells through tissue-tissue interactions during tongue morphogenesis. Dev Biol 341, 186–95 (2010).

28. Ishan, M. et al. Increased activity of mesenchymal ALK2-BMP signaling causes posteriorly truncated microglossia and disorganization of lingual tissues. Genesis 58, e23337 (2020).

29. Iwata, J., Suzuki, A., Pelikan, R.C., Ho, T.V. & Chai, Y. Noncanonical transforming growth factor β (TGFβ) signaling in cranial neural crest cells causes tongue muscle developmental defects. J Biol Chem 288, 29760–70 (2013).

30. Jeong, J., Mao, J., Tenzen, T., Kottmann, A.H. & McMahon, A.P. Hedgehog signaling in the neural crest cells regulates the patterning and growth of facial primordia. Genes Dev 18, 937–51 (2004).

31. Jonker, L., Kist, R., Aw, A., Wappler, I. & Peters, H. Pax9 is required for filiform papilla development and suppresses skin-specific differentiation of the mammalian tongue epithelium. Mech Dev 121, 1313–22 (2004).

32. Kajimura, S., Seale, P. & Spiegelman, B.M. Transcriptional control of brown fat development. Cell Metab 11, 257–62 (2010).

33. Lan, Y., Wang, Q., Ovitt, C.E. & Jiang, R. A unique mouse strain expressing Cre recombinase for tissue-specific analysis of gene function in palate and kidney development. Genesis 45, 618–24 (2007).

34. Li, J., Mao, C., Ma, L. & Zhou, X. Giant sublingual hamartoma with medial cleft tongue: a case report and literature review. J Int Med Res 48, 300060520942089 (2020).

35. Liu, H.X. et al. Separate and distinctive roles for Wnt5a in tongue, lingual tissue and taste papilla development. Dev Biol 361, 39–56 (2012).

36. Macca, M. & Franco, B. The molecular basis of oral-facial-digital syndrome, type 1. Am J Med Genet C Semin Med Genet 151C, 318–25 (2009).

37. Manjila, S. et al. Duplication of the pituitary gland associated with multiple blastogenesis defects: Duplication of the pituitary gland (DPG)-plus syndrome. Case report and review of literature. Surg Neurol Int 3, 23 (2012).

38. Martín, L.P. et al. Atypical case of congenital maxillomandibular fusion with duplication of the craniofacial midline. Craniomaxillofac Trauma Reconstr 4, 113–20 (2011).

39. Millington, G. et al. Cilia-dependent GLI processing in neural crest cells is required for tongue development. Dev Biol 424, 124–137 (2017).

40. Mistretta, C.M. & Liu, H.X. Development of fungiform papillae: patterned lingual gustatory organs. Arch Histol Cytol 69, 199–208 (2006).

41. Morita, H. et al. Neonatal lethality of LGR5 null mice is associated with ankyloglossia and gastrointestinal distension. Mol Cell Biol 24, 9736–43 (2004).

42. Mossaad, A.M., Abdelrahman, M.A., Ibrahim, M.A. & Al Ahmady, H.H. Surgical Management of Facial Features of Robinow Syndrome: A Case Report. Open Access Maced J Med Sci 6, 536–539 (2018).

43. Mostafa, M.I., Temtamy, S.A., el-Gammal, M.A. & Mazen, I.M. Unusual pattern of inheritance and orodental changes in the Ellis-van Creveld syndrome. Genet Couns 16, 75–83 (2005).

44. Murcia, N.S. et al. The Oak Ridge Polycystic Kidney (orpk) disease gene is required for left-right axis determination. Development 127, 2347–55 (2000).

45. Nguyen, A.P., Firth, N., Mougos, S. & Kujan, O. Lingual Leiomyomatous Hamartoma in an Adult Male. Case Rep Dent 2018, 4162436 (2018).

46. Obregon, M.J. Adipose tissues and thyroid hormones. Front Physiol 5, 479 (2014).

47. Ohazama, A. et al. Lrp4 modulates extracellular integration of cell signaling pathways in development. PLoS One 3, e4092 (2008).

48. Okuhara, S. et al. Temporospatial sonic hedgehog signalling is essential for neural crest-dependent patterning of the intrinsic tongue musculature. Development 146(2019).

49. Parada, C., Han, D. & Chai, Y. Molecular and cellular regulatory mechanisms of tongue myogenesis. J Dent Res 91, 528–35 (2012).

50. Rao, A.Y. Complete Midline Cleft of Lower Lip, Mandible, Tongue, Floor of Mouth with Neck Contracture: A Case Report and Review of Literature. Craniomaxillofac Trauma Reconstr 8, 363–9 (2015).

51. Pauws, E. et al. Tbx22null mice have a submucous cleft palate due to reduced palatal bone formation and also display ankyloglossia and choanal atresia phenotypes. Hum Mol Genet 18, 4171–9 (2009).

52. Rother, P., Wohlgemuth, B., Wolff, W. & Rebentrost, I. Morphometrically observable aging changes in the human tongue. Ann Anat 184, 159–64 (2002).

53. Sambasivan, R., Kuratani, S. & Tajbakhsh, S. An eye on the head: the development and evolution of craniofacial muscles. Development 138, 2401–15 (2011).

54. Scarda, A. et al. Increased adipogenic conversion of muscle satellite cells in obese Zucker rats. Int J Obes (Lond) 34, 1319–27 (2010).

55. Seale, P. et al. PRDM16 controls a brown fat/skeletal muscle switch. Nature 454, 961–7 (2008).

56. Senan, M. & Menon, V.P. Pentafid tongue: A new entity. Indian J Plast Surg 48, 301–4 (2015).

57. Sidossis, L. & Kajimura, S. Brown and beige fat in humans: thermogenic adipocytes that control energy and glucose homeostasis. J Clin Invest 125, 478–86 (2015).

58. Soriano, P. Generalized lacZ expression with the ROSA26 Cre reporter strain. Nat Genet 21, 70–1 (1999).

59. Srinivas, S. et al. Cre reporter strains produced by targeted insertion of EYFP and ECFP into the ROSA26 locus. BMC Dev Biol 1, 4 (2001).

60. Teo, D.T., Johnson, R.F. & McClay, J.E. An unusual tongue base mass in an infant: Tongue base sialolipoma. Ear Nose Throat J 94, E14–6 (2015).

61. van Bokhoven, H. et al. Mutation of the gene encoding the ROR2 tyrosine kinase causes autosomal recessive Robinow syndrome. Nat Genet 25, 423–6 (2000).

62. Vettor, R. et al. The origin of intermuscular adipose tissue and its pathophysiological implications. Am J Physiol Endocrinol Metab 297, E987–98 (2009).

63. Waters, D.L. Intermuscular Adipose Tissue: A Brief Review of Etiology, Association With Physical Function and Weight Loss in Older Adults. Ann Geriatr Med Res 23, 3–8 (2019).

64. Xu, J. et al. Hedgehog signaling patterns the oral-aboral axis of the mandibular arch. Elife 8(2019).

65. Yi, R. et al. Morphogenesis in skin is governed by discrete sets of differentially expressed microRNAs. Nat Genet 38, 356–62 (2006).

66. Yin, S. & Zhao, Z. Incomplete Cleft Palate, Bifid Tongue, and Oral Hamartomas: A Rare Congenital Anomaly. J Craniofac Surg 31, e184–e185 (2020).

67. Yuan, G. et al. Cleft Palate and Aglossia Result From Perturbations in Wnt and Hedgehog Signaling. Cleft Palate Craniofac J 54, 269–280 (2017).

68. Zaghloul, N.A. & Brugmann, S.A. The emerging face of primary cilia. Genesis 49, 231–46 (2011).

69. Zhong, Z., Zhao, H., Mayo, J. & Chai, Y. Different requirements for Wnt signaling in tongue myogenic subpopulations. J Dent Res 94, 421–9 (2015).

70. Ziermann, J.M., Diogo, R. & Noden, D.M. Neural crest and the patterning of vertebrate craniofacial muscles. Genesis 56, e23097 (2018).

